# The HDAC inhibitor CI-994 acts as a molecular memory aid by facilitating synaptic and intra-cellular communication after learning

**DOI:** 10.1101/2021.09.21.460970

**Authors:** Allison M Burns, Mélissa Farinelli-Scharly, Sandrine Hugues-Ascery, Jose Vicente Sanchez-Mut, Giulia Santoni, Johannes Gräff

## Abstract

Long-term memory formation relies on synaptic plasticity, activity-dependent transcription and epigenetic modifications. Multiple studies have shown that HDAC inhibitor (HDACi) treatments can enhance individual aspects of these processes, and thereby act as putative cognitive enhancers. However, their mode of action is not fully understood. In particular, it is unclear how systemic application of HDACis, which are devoid of substrate specificity, can target pathways that promote memory formation. In this study, we explore the electrophysiological, transcriptional and epigenetic responses that are induced by CI-994, a class I HDAC inhibitor, combined with contextual fear conditioning (CFC) in mice. We show that CI-994-mediated improvement of memory formation is accompanied by enhanced long-term potentiation in the hippocampus, a brain region recruited by CFC, but not in the striatum, a brain region not primarily implicated in contextual memory formation. Furthermore, using a combination of bulk and single cell RNA sequencing, we find that synaptic plasticity-promoting gene expression cascades are more strongly engaged in the hippocampus than in the striatum, but only when HDACi treatment co-occurred with CFC, and not by either treatment alone. Lastly, using ChIP-sequencing, we show that the combined action of HDACi application and conditioning is required to elicit enhancer histone acetylation in pathways that may underlie improved memory performance. Together, our results indicate that systemic HDACi administration amplifies brain-region specific processes that are naturally induced by learning. These findings shed light onto the mode of action of HDACis as cognitive enhancers.

**Significance Statement:** Memory formation relies on a plethora of functions, including epigenetic modifications. Over the past years, multiple studies have indicated the potential of HDAC inhibitors (HDACi) to act as cognitive enhancers, but their mode of action is not fully understood. Here, we tested whether HDACi treatment improves memory formation via “cognitive epigenetic priming”, stipulating that HDACis – without inherent target specificity – specifically enhance plasticity-related processes. We found that combining HDACi with fear learning, but not either treatment alone, enhances synaptic plasticity as well as memory-promoting transcriptional signaling in the hippocampus, a brain area known to be recruited by fear learning, but not in others. These results lend experimental support to the theory of “cognitive epigenetic priming”.

## Introduction

Long-term memory is a product of synaptic communication as well as activity-dependent transcription that is regulated by epigenetic signaling (1–5). For example, memory forming tasks, such as contextual fear conditioning (CFC), are paralleled by gene expression and histone acetylation changes in the hippocampus (6–8), while impaired cognition, as seen in Alzheimer’s Disease and age-related cognitive decline, is coupled with a reduction in hippocampal histone acetylation and plasticity-related gene expression (9–13). Some of these epigenetic and transcriptional changes can be augmented by systemic HDAC inhibitor (HDACi) treatment, which improves memory in both healthy and cognitively impaired mice (10–12, 14, 15). Although the use of HDACis in these studies testifies to their suitability as pharmacological memory aids, the mechanisms by which HDACi enhances memory are not fully understood. In particular, it is unclear how systemic application of HDACis, most of which are devoid of substrate specificity *per se*, can target pathways that promote memory formation.

One proposed theoretical mode of action for HDACis as cognitive enhancers is called “cognitive epigenetic priming” (3, 16). This model is inspired by evidence from cancer research, where several HDACis have been shown to improve target efficacy of anti-cancer treatments (17, 18), and from addiction research, where chronic drug abuse was found to durably enrich histone acetylation, which relaxes the chromatin structure into a primed state and thereby lowers the activation threshold for gene expression changes during subsequent drug exposures (19, 20). Analogously, for cognition, this theory stipulates that by broadly increasing histone acetylation, HDACi treatment leads to an overall primed state. Memory-induced neuronal activity, which is inherently characterized by a high degree of target specificity (2), would then further enrich HDACi-induced histone acetylation and recruit the transcriptional machinery specifically to synaptic plasticity-related genes.

In this study, we tested the concept of “cognitive epigenetic priming” in mice on three different levels. First, we investigated whether systemic HDACi treatment elicits brain-region specific electrophysiological and transcriptional responses after contextual fear conditioning, a hippocampus-dependent memory task. Second, we assessed whether and to which extent specific cell types are affected by the HDACi treatment in combination with learning using single nuclear RNA-sequencing (snRNA-seq) of the hippocampus; and third, we determined which gene loci are epigenetically regulated by HDACi treatment using chromatin immunoprecipitation (ChIP) followed by sequencing. These experiments were designed to better understand the underlying mechanisms of HDACis as potential cognitive enhancers.

## Results

### Systemic HDACi treatment enhances memory consolidation after subthreshold contextual fear conditioning

To investigate the mechanisms by which systemic HDACi treatment enhances fear memory, we treated mice with the HDACi CI-994, before subjecting them to a subthreshold contextual fear conditioning (CFC) task, a modified Pavlovian conditioning paradigm that, alone, does not induce memory formation (21). CI-994 is a class I HDACi that selectively impedes HDACs 1-3 (22), promotes functional recovery after stroke (23), and that has shown promise against cognitive dysfunctions in preclinical animal models (15, 24, 25). When i.p. injected it crosses the blood-brain-barrier and remains in the brain at concentrations greater than 1000nm for up to 5 hours (15). One hour prior to CFC or Context only exposure (Context), mice were interperitoneally (i.p.) injected with 30mg/kg of CI-994 or its vehicle (VEH) (**Fig. 1A**). One day later, freezing was measured during a 3 min context exposure. We found that pairing the subthreshold CFC paradigm with the HDACi significantly improved memory retention (*P* = 0.0002) compared to the CFC paradigm alone (*P* = 0.172), and compared to HDACi treatment alone (*P* = 0.997, Tukey’s HSD test following one-way ANOVA, F_(3,39)_ = 10.16, *P* = 4.44e-05) (**Fig. 1B**). There were no freezing differences between context and CFC exposure for VEH-treated animals (*P* = 0.172). Furthermore, HDACi treatment had no effect on speed (F_(3,39)_ = 1.71, *P* = 0.18) or distance travelled (F_(3,39)_ = 1.69, *P* = 0.184) and did not affect anxiety levels as measured by an open field test at the time of initial encoding (F_(3,39)_ = 0.536, *P* = 0.66) (**Supplemental Fig. 1**). These results indicate that the HDACi treatment can elevate an otherwise inefficient learning paradigm above threshold and lead to long-term memory retention.

**Figure 1.**
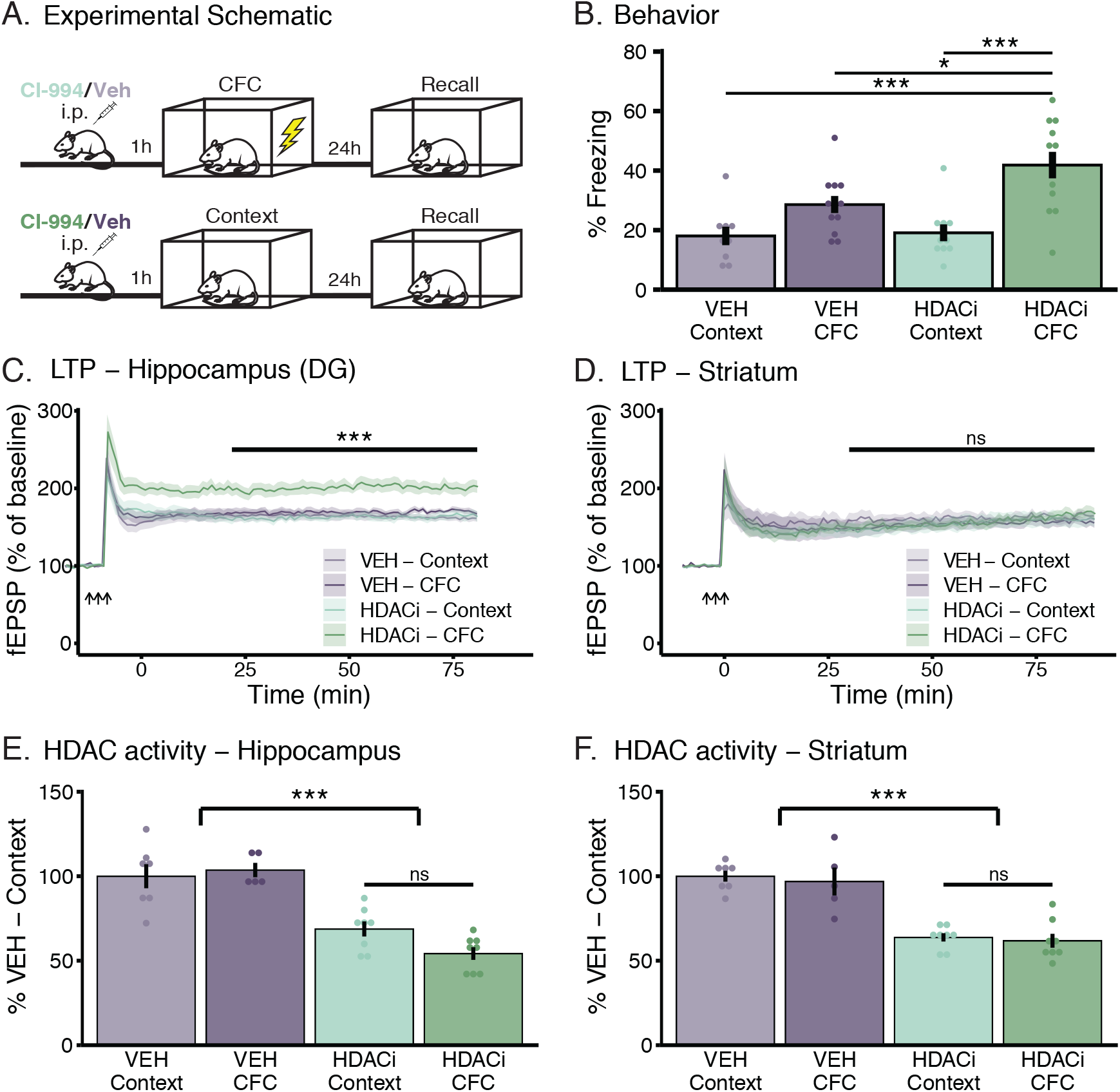
HDACi treatment enhances long term potentiation in the hippocampus, but not the striatum, despite reducing HDAC activity in both brain regions. **(A)** Schematic representation of the behavioral paradigm for subthreshold contextual fear conditioning (CFC) (top) and Context only exposure (bottom). All animals received an i.p. injection of either vehicle or of the HDACi, CI-994 (30mg/kg). One hour later, animals underwent sub-threshold CFC (2x 0.2mA – 1s shocks) and fear memory was measured one day later by re-exposing animals to the conditioning context in the absence of the foot shock. **(B)** HDACi combined with CFC increases the percent of time spent freezing (> 1s) during 3-minute re-exposure to the conditioning chamber. n= 9-12 animals/group. **(C and D)** HDACi combined with CFC enhances LTP in response to 3 trains of high frequency stimulation (HFS – arrows) in the perforent pathway of the dentate gyrus **(C)** but not in the corti-cal-striatal pathway **(D)** one hour after conditioning. Statistical differences were calculated for the 30 minutes (end of short-term-potentiation) to 90 minutes (end of recording) for each mouse. n = 8 animals/group. **(E and F)** HDAC activity was reduced after HDACi in both the hippocampus **(E)** and striatum **(F)** with no further reduction in HDAC activity in response to CFC. One or two-way ANOVA with Tukey’s HSD multiple-comparisons test was used for analysis. Graphs represent mean + SEM. * *P* < 0.05, ** *P* < 0.01, *** *P* < 0.001

### Systemic HDACi treatment regulates long term potentiation in an activity-specific manner

To explore whether HDACi treatment improves memory via cognitive epigenetic priming, we first assessed its mode of action on synaptic plasticity. To this end, we measured the effects of HDACi on long term potentiation (LTP) in the hippocampus, a brain region activated by CFC (26), and the striatum, a brain region that is not directly involved in contextual memory formation (27) one hour after CFC. We found a significant increase in LTP at perforant path synapses of the dentate gyrus (DG) of the hippocampus when CFC was paired with the HDACi treatment (**Fig. 1C**; one-way ANOVA, F_(3,28)_ = 10.57, *P* = 8.09e-05). Without CFC, the HDACi had no effect on LTP; similarly, CFC alone did not facilitate LTP. Conversely, at cortico-striatal fibers, the HDACi treatment had no effect regardless of the behavioral paradigm (F_(3,28)_ = 0.234, p = 0.872) (**Fig. 1D**). Neither paired pulse facilitation (PPF) nor input/output (I/O) relationships were changed in either brain region (**Supplemental Fig. 2A-D**). Importantly, combining CFC with HDACi also enhanced LTP at Schaffer collaterals of the CA1, another hippocampal subregion (one-way ANOVA, F_(3,28)_ = 5.213, *P* = 0.005) both after sub-threshold CFC (**Supplemental Fig. 3**), and when using a stronger CFC paradigm (one-way ANOVA, F_(3,33)_ = 3.663, *P* = 0.0221) (**Supplemental Fig. 4**).

These findings indicate a brain area-specific effect of the HDACi treatment, with only brain areas engaged by CFC displaying enhanced synaptic plasticity. Interestingly, this brain region-specific effect on synaptic plasticity occurred in spite of the same degree of HDAC activity inhibition in both brain regions. HDAC activity was reduced by about 50% in both the hippocampus (F_(1,24)_ = 60.15, p = 5.44e-08) (**Fig. 1E**) and the striatum (F_(1,24)_ = 68.96, p = 1.62e-08) (**Fig. 1F**) in response to HDACi, with no difference in HDAC activity induced by learning itself. Thus, despite the same extent of HDAC inhibition induced by the HDACi, synaptic plasticity was only altered in the brain area directly engaged by CFC.

To confirm these findings in a different task, and to show that the HDACi does not only improve plasticity and performance in a hippocampus-specific manner, we also tested HDACi treatment during rotarod training, a motor skill learning task known to depend on the cortico-striatal pathway (28). Animals were i.p. injected with HDACi or VEH one hour before training (**Supplemental Fig. 5A**). We found that HDACi-treated animals were able to stay on the apparatus for longer than their VEH-treated counterparts (one-way ANOVA, F_(1,120)_ =12.155, p = 0.0007) (**Supplemental Fig. 5B**), indicating improved motor learning. While neither training nor HDACi had any effect on hippocampal or striatal LTP (**Supplemental Fig. 5C-D**), we found that HDACi paired with rotarod training selectively increased striatal PPF (two-way ANOVA, F_(3,192)_ = 12.217, *P* = 2.37e-07) (**Supplemental Fig. 5E-F**), which is known to underlie motor learning in the striatum (29). In addition, there were no major differences in I/O in the striatum or the hippocampus (**Supplemental Fig. 5G-H**). These electrophysiological data are thus in support of the cognitive epigenetic priming hypothesis at the level of these two brain areas, insofar as the HDACi application *per se* did not yield any measurable difference, but necessitated task-specific neuronal activity to reveal its potentiating effect.

### HDACi activates different transcriptional cascades in response to CFC in the hippocampus and striatum

To further understand the molecular mechanisms by which epigenetic priming leads to improved memory performance, we used bulk RNA-sequencing in the hippocampus and striatum to determine which genes are activated when CFC is combined with HDACi treatment. For this, we extracted and sequenced total mRNA from whole-tissue homogenates one hour after CFC or context only exposure, using the same experimental setup as for the electrophysiological recordings (**Fig. 1A**). The Illumina HiSeq4000 was used to generate four replicate libraries for each group with a minimum of 28M uniquely mapping paired reads per sample (**Supplemental Fig. 6A**). In total, 26,020 genes were expressed by greater than 1 count per million (CPM) in at least 4 of the libraries. Principal component analysis for the top 1000 most variable genes across all libraries revealed that 93% of the variance results from inter-brain region differences (**Supplemental Fig. 6B, C**).

In the hippocampus, consistent with previous data (30), we found no differentially expressed genes (DEGs) (P ≤ 0.05; |log_2_FC| ≥ 0.4) between CFC and context only exposure in VEH-treated animals (**Fig. 2A, left panel**). Likewise, when comparing CFC with the context-only group in HDACi-treated animals, no DEGs were detected, indicating that subthreshold CFC alone is not sufficient to induce detectable transcriptional changes(**Fig. 2A, middle left panel**). Conversely, when context exposure was paired with CI-994, we found 1002 and 1679 genes significantly up- and downregulated, respectively, indicating that the addition of the HDACi alone alters the transcriptional landscape (**Fig. 2A, middle right panel**). When HDACi-CFC was compared to VEH-CFC, we detected 1336 up-regulated genes, a 25% increase when compared to the context only contrast, but a similar number (1608) of downregulated genes (**Fig. 2A, right panel**).

**Figure 2.**
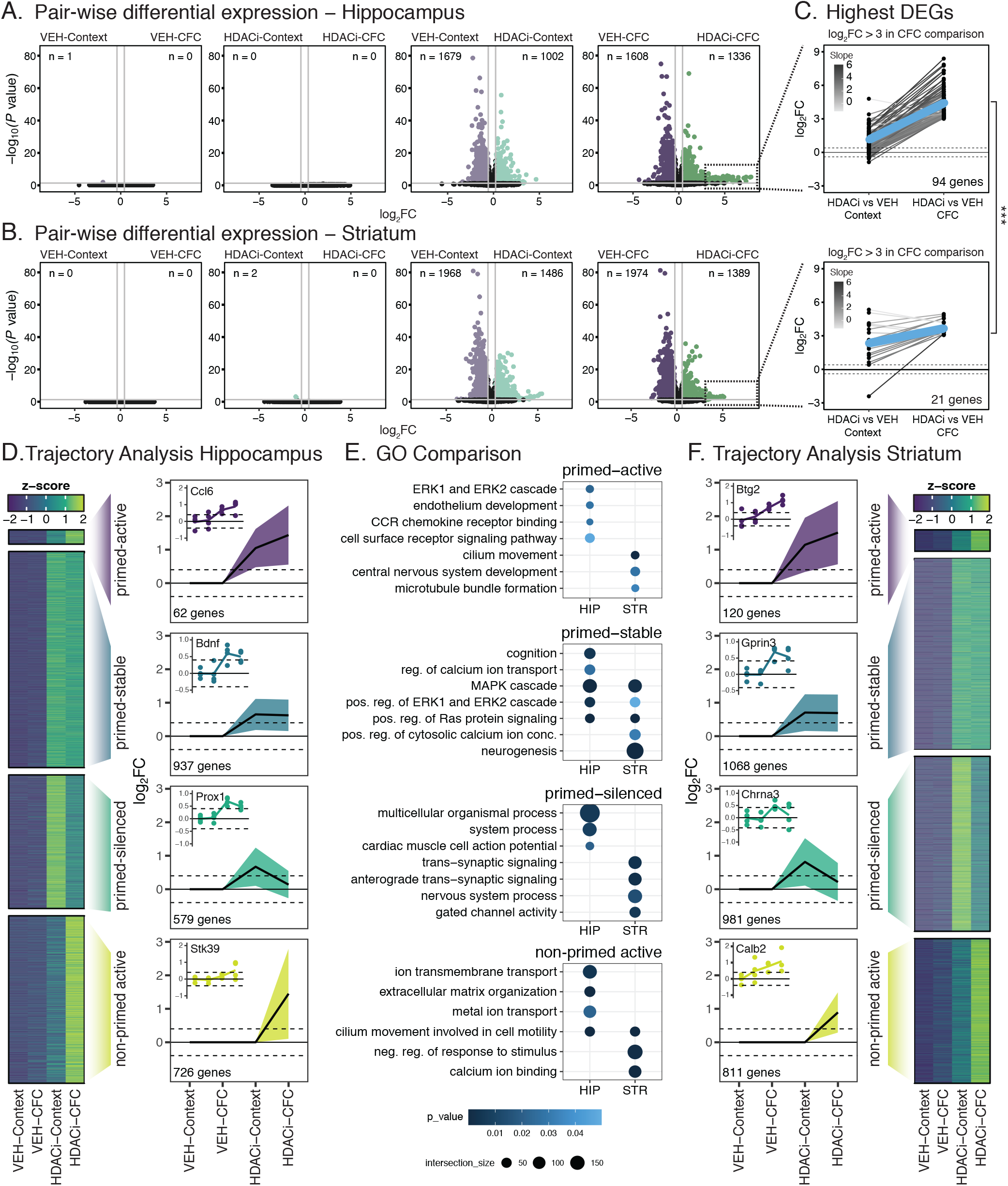
HDACi activates brain region-specific transcriptional cascades in response to CFC. **(A and B)**Volcano plots of magnitude of differential expression (log_2_FC) versus statistical significance (-log_10_ *P*-value) of pairwise comparisons labelled above each plot in the hippocampus (**A**) and striatum (**B**). n-values in corners represent number of DEGs (log_2_FC ≥ 0.4; adjusted *P*-value ≤ 0.05) in the corresponding corner label. **(C)** Comparisons of DEGs that have ≥ 3 log_2_FC in the HDACi-CFC compared to VEH-CFC (right column) in the hippocampus (top) and striatum (bottom). Log_2_FC values are plotted for those same genes in the HDACi-Context compared to VEH-Context in the left column. Lines connect the same gene in each comparison and are colored by log_2_FC difference between the two comparisons (slope). The blue line represents the average slope for each brain region. Student’s t-test comparing slopes between hippocampus and striatum (right of plots). **(D)** Heat map of z-scores of average gene counts in the hippocampus (left). Line graphs in trajectory plots represent significant log_2_FC values for each group (right). Count in lower left corner indicates number of genes in each cluster. Line plots shown as mean ±SEM. Insets represent DEGs from each cluster. Normalized counts for each replicate were compared to average normalized count for VEH-Context. **(E)** Gene ontology (GO) analysis of hippocampal (left) and striatal (right). **(F)** Gene cluster analysis for striatum RNA-seq data as in D. n = 4 biologically independent samples. * *P* < 0.05, ** *P* < 0.01, *** *P* < 0.001

In the striatum, there were no DEGs between the CFC and context only exposure in either the VEH or HDACi treated animals (**Fig. 2B, left panels**). Similar to the hippocampus, when HDACi-Context was compared to VEH-Context, 1486 and 1968 genes were significantly up- and downregulated (**Fig. 2B, middle right panel**). In contrast to the hippocampus, however, no further increase in the number of DEGs was detected when HDACi was paired with CFC (1389 genes were upregulated, and 1974 genes were downregulated) (**Fig. 2B, right panel**). Lists of pair-wise differential expression for both the hippocampus and striatum can be found in **Supplemental Table 1**.

When focusing on strongly up-regulated genes (P ≤ 0.05, log_2_FC ≥ 3), we detected 4.5x more genes in the hippocampus than in the striatum (**Fig. 2C**). Furthermore, these genes were more strongly activated in the hippocampus than in the striatum (Student’s t-test of slope values, P = 2.304e-05) (**Fig. 2C**). The strongly upregulated hippocampal genes (**Supplemental Table 2**) were enriched in ion-transport ontologies and included transthyretin, *Ttr*, which has been shown to provide neuroprotection in aged mice and to be associated with enhanced memory (31, 32), and synaptotagmin 13 (*Syt13*), a gene previously shown to be up-regulated after CFC (33). In contrast, the striatal genes were primarily predicted genes (**Supplemental Table 2**) and GO analysis did not yield any enriched pathways. This indicates that in the hippocampus, the expression of these genes is further enhanced when the HDACi is paired with CFC, while pairing HDACi with CFC had no such effect in the striatum.

Next, we set out to identify transcriptional patterns by selecting genes that were differentially expressed (*P* <= 0.05, log_2_FC >= 0.4) when compared to the baseline group (VEH-Context). For this, all DEGs underwent decision tree clustering as described in the materials and methods. Considering that we aim to specifically understand the targets of epigenetic priming, we focused on genes that are up-regulated by HDACi treatment in our analyses, however the other clusters, including down-regulated ones, as well as their associated genes, can be found in **Supplemental Fig. 7** and **Supplemental Table 3**. Four major clusters were identified as trajectories of interest (**Fig. 2D**). In the hippocampus, we found 1) 62 genes that were up-regulated by the HDACi treatment alone (i.e., in the HDACi-VEH group) and further increased when HDACi was paired with CFC (i.e, the HDACi-CFC group), which we termed *primed-active*; 2) 937 genes that were increased by HDACi treatment but showed no further CFC-driven increase, which we termed *primed-stable*; 3) 579 genes that were enriched by HDACi treatment, but were reduced when the HDACi was paired with CFC, which we termed *primed-silenced*; 4) and 726 genes that were only activated when combining HDACi with CFC, but not by either condition alone, which we termed *non-primed active* (**Fig. 2D**). In the striatum, the order of magnitude of DEGs was similar (**Fig. 2F**). There were 120 genes in the *primed-active* cluster, 1068 genes in the *primed-stable* cluster, 981 genes in the *primed-silenced* cluster and 811 genes in the *non-primed active cluster* (**Fig. 2F**).

We next performed a Gene Ontology (GO) analysis (**Fig. 2E**) to identify enriched pathways in each cluster in both the hippocampus and the striatum. In the hippocampus, the *primed-active* cluster was enriched for the Erk1 and Erk2 cascade, which has been implicated in synaptic plasticity as well as learning and memory (34, 35). This cluster included cytokine genes, such as *Ccl6* (**Fig. 2D**), which is involved in the p38-MAPK pathway (36, 37) and which plays a role in cell survival (38). Conversely, in the striatum, the *primed-active* cluster was not enriched for any ontologies involved in MAPK/ERK signaling or learning and memory.

Furthermore, the *primed-stable* cluster was characterized by learning and memory-related pathways such as cognition and regulation of calcium ion transport in the hippocampus, but not the striatum (**Fig. 2E**). Hippocampal DEGs in this cluster included brain derived neurotrophic factor (*Bdnf*) and the proto-oncogene Jun (*Jun*), both immediate early genes (IEGs) induced by neuronal activity and implicated in synaptic plasticity as well as learning and memory. *Bdnf* plays a critical role in hippocampal CFC (39) and enhances synaptic strength at the Schaffer collateral-CA1 synapses (40), while *Jun* is a member of the AP-1 transcriptional activator complex, which binds enhancers and regulates chromatin opening during CFC (41, 42). In the striatum, this cluster did not include memory-related IEGs. It did, however, contain pathways involved in intracellular signal transduction that also regulate learning and memory (6, 43) such as the “MAPK cascade”, “Ras protein signal transduction” and “Erk1 and Erk2 cascade”. This comparison stipulates that HDACi similarly primes the MAPK pathway in both the hippocampus and the striatum but further potentiates only the *primed-active* genes in the hippocampus.

In the *primed-silenced* or *non-primed active* states, no ontologies associated with synaptic signal transduction were found (**Fig. 2E**). Finally, the hippocampal *non-primed active* cluster, represented by genes that are only transcribed after combined HDACi-CFC, is enriched for “metal ion transport” and “ion transmembrane transport” pathways, while in the striatum, it is enriched for genes involved in a “negative response to stimulus”. This could indicate that the combination of HDACi treatment and CFC increases inhibitory signalling in the striatum, possibly related to the decreased motor response following conditioning. Of note, none of the clusters in which HDACi reduced transcription included pathways that are involved in learning and memory or synaptic plasticity in either the hippocampus or the striatum. Taken together, these results illustrate that HDACi treatment, and not CFC, is the major driver of differential transcription between the hippocampus and the striatum. It enhances the Mapk/Erk signalling pathway in both the hippocampus and the striatum as seen in the comparisons of the *primed-stable* groups, but is able to further induce learning-specific genes in the *primed-active* state in the hippocampus when paired with CFC, but not the striatum.

### HDACi activates different transcriptional cascades across cell types within the hippocampus

Next, we aimed to understand which cell types within the hippocampus are most responsive to HDACi treatment. To do so, we used snRNA-seq on isolated hippocampi from animals that were treated with either HDACi or VEH one hour before undergoing CFC. Since transcriptional differences were most prominent in the HDACi-CFC versus the VEH-CFC groups (**Fig. 2C**), we focused on only this comparison. We performed dimensionality reduction using uniform manifold approximation and projection (UMAP) and clustered nuclei by the k-nearest neighbors. We removed clusters containing fewer than 50 nuclei, revealing 30 distinct clusters consisting of 15,339 total nuclei and expressing a total of 24,271 genes (**Supplemental Fig. 8A**). These clusters were then assigned to known cell types by comparing expression of cell-type specific genes taken from previously published snRNA-seq datasets (44–47) and the Allen Brain Atlas (48) (**Supplemental Fig. 8B**). This analysis identified 10 distinct cell types: 4 clusters of excitatory neurons that split based on location within the hippocampus (5175 DG nuclei, 2871 CA1 nuclei, 1657 CA3 nuclei, and 507 nuclei with no location marker); 1 cluster of 794 inhibitory neurons; 4 glial clusters (1960 Oligodendrocytes, 254 oligo-precursors, 763 astrocytes and 880 microglia); and a final cluster of 478 nuclei (NA) which could not be assigned to a single cell type based on its expression profile (**Fig. 3A and Supplemental Fig. 8C**). In line with previous work (49), neuronal clusters had more expressed genes than glial clusters (**Supplemental Fig. 8D**) and the proportions of cell types were similar to those reported for the hippocampal region in the Blue Brain Atlas (50) (**Supplemental Fig. 8E**).

**Figure 3.**
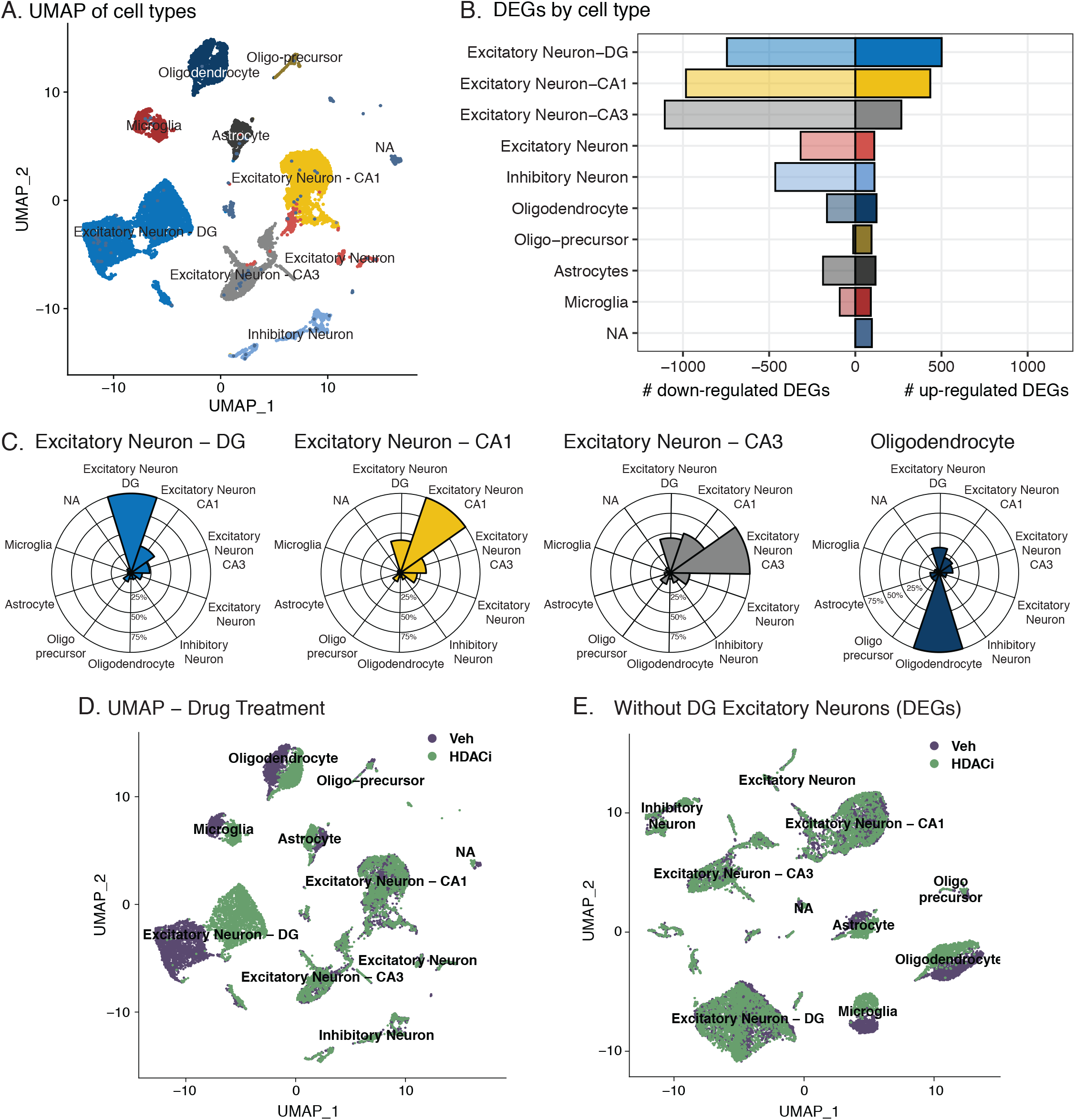
HDACi activates different transcriptional cascades within hippocampal cell types. **(A)** Uniform manifold approximation and projection (UMAP) visualization of 15,339 nuclei from the full hippocampus colored by 10 identified cell-types. NA refers to nuclei that could not be assigned a cell type based on expression of marker genes. **(B)** Number of up-regulated (right; log_2_FC ≥ 1; FDR ≤ 0.05) and down-regulated (left; log_2_FC ≤ −1; FDR ≤ 0.05) genes in each cell type when comparing HDACi-CFC to VEH-CFC. **(C)** Radar plots showing overlap of up-regulated genes across cell types. (Left) Percent overlap of Excitatory Neurons - DG with others. (Middle left) Percent overlap of Excitatory Neurons – CA1 with other clusters. (Middle right) Percent overlap of Excitatory Neurons – CA3 with other clusters. (Right) Percent overlap of Oligodendrocytes with other clusters. **(D)** UMAP visualization of nuclei from the full hippocampus colored by sample drug treatment. **(E)** UMAP visualization, colored by drug treatment, after removing the 501 up-regulated genes in the DG excitatory neurons and re-clustering. n = 2 biological replicates per group (HDACi-CFC and VEH-CFC).

We then explored whether pairing CFC with HDACi induces distinct responses across cell types. Augur, a tool that prioritizes a population’s responsiveness to an experimental perturbation (51), reported a similar global responsiveness for all clusters (**Supplemental Fig. 9A**), and with the exception of oligo-precursors, HDACi treatment did not significantly change cell type composition within clusters (**Supplemental Fig. 9B**). However, HDACi treatment differentially regulated a distinct set of genes in each cell type, with excitatory neurons having the largest HDACi response (**Fig. 3B, Supplemental Fig. 9C** and **Supplemental Table 4**). These DEGs were highly cluster specific: We found that excitatory neurons of the DG share 36% and 24% of their up-regulated DEGs with excitatory neurons of the CA1 and CA3, respectively, and fewer than 15% with each of the other cell types (**Fig. 3C, left panel**). This low overlap of HDACi induced up-regulation was also seen in other cell types (**Fig. 3C** and **Supplemental Fig. 9D**). Among excitatory neurons, 45%, 38% and 27% of genes were uniquely up-regulated in the DG, CA1 and CA3, respectively, while among glial cells, 68%, 53% and 47% were uniquely upregulated among microglia, astrocytes and oligodendrocytes, respectively (**Supplemental Fig. 9E)**. Down-regulated genes also appeared to be cluster specific, although to a lower magnitude than the up-regulated genes (**Supplemental Fig. 9F**).

Interestingly, we found an HDACi-specific separation for excitatory neurons in the DG and for glia, but not for any other cluster (**Fig. 3D**). This split was mainly mediated by the upregulated genes within the DG, as removing those genes and re-running the dimension reduction re-merged the split DG cluster (**Fig. 3E**). Conversely, there was no cluster re-merging when removing up-regulated DEGs from CA1, glia or from any other cell type (**Fig. 3F and Supplemental Fig. 10A**). Furthermore, DG cluster re-merging was specific to the up-regulated genes, as removing only downregulated genes had no effect (**Supplemental Fig. 10B**). Together, these results provide supporting evidence that pairing CFC with HDACi treatment transcriptionally activates different gene sets across cell types, with a particularly strong response among upregulated genes in the DG. For this reason, we continued our analysis of epigenetic priming by focusing on excitatory neurons of the DG.

### HDACi combined with CFC enriches H3K27ac at genes involved in synaptic communication

Given the strong up-regulation of genes involved in excitatory neurons of the DG, we characterized histone acetylation in this region by chromatin immunoprecipitation followed by sequencing (ChIP-Seq). We focused on H3K27ac, a known marker of active enhancers that is enriched at activity-dependent regulatory elements after neuronal activation (30, 42, 52–54), correlates with gene transcription (30, 55) and often co-occurs with H3K9ac, a marker of active promoters (56). Furthermore, in line with previous studies (15, 25, 57, 58), we found that HDACi treatment increased global H3K27ac, alongside H3K9ac and H4K12ac as revealed by western blotting (Two-way ANOVA, F_(3,60)_ = 22.47, *P* = 1.11e-13) (**Supplemental Fig. 11**). For ChIP-seq, we had 3 replicates, each from the pooled DG from 5 mice and sorted NeuN+ nuclei by fluorescence activated nuclear sorting (FANS) (**Fig. 4A**). Libraries for the H3K27ac-immunoprecipitated samples were prepared and processed as described in the materials and methods.

**Figure 4.**
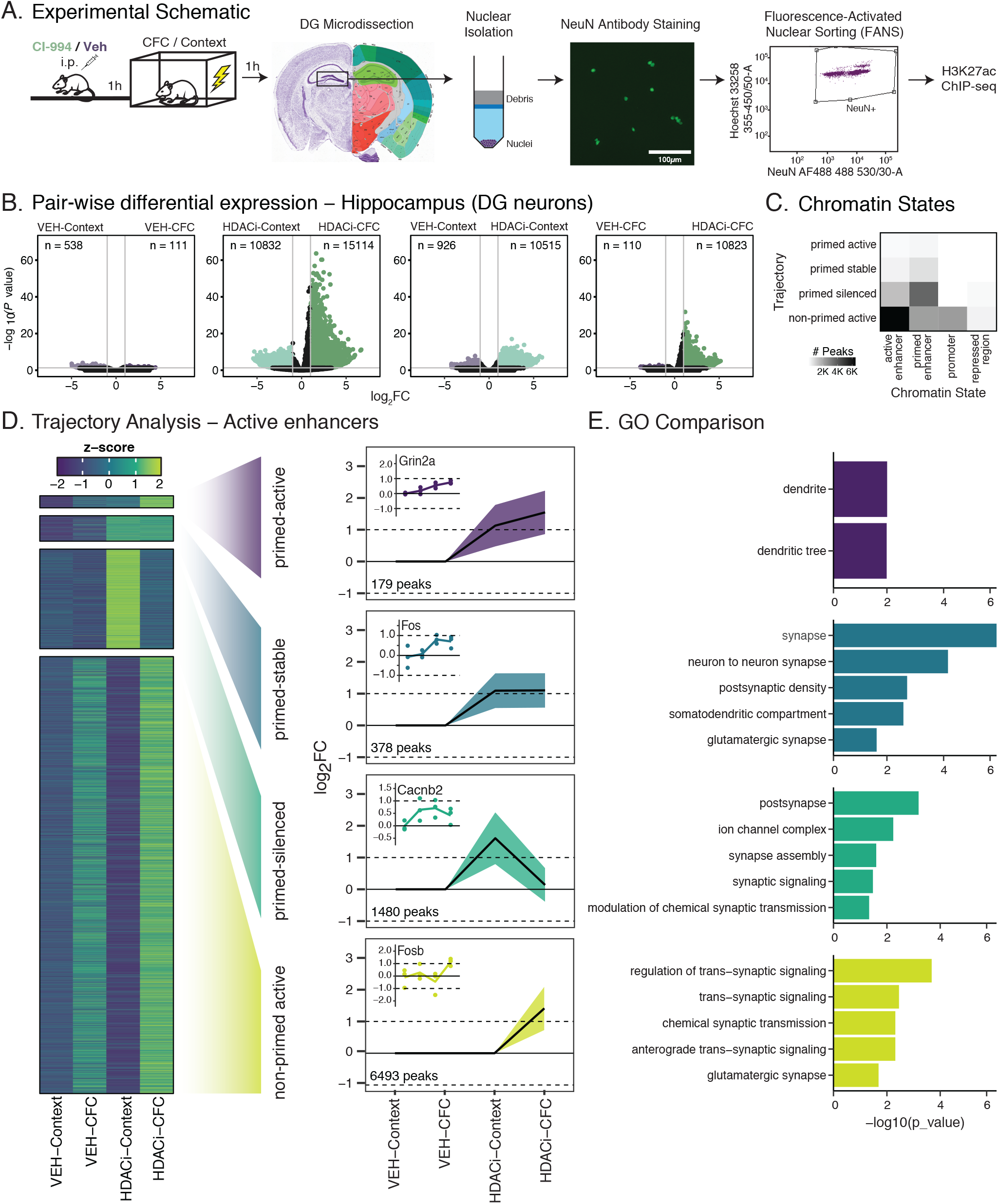
HDACi enriches H3K27ac at genes involved in neuronal synaptic communication. **(A)** Schematic of experimental outline. **(B)** Volcano plots showing the magnitude of differential H3K27ac enrichment (log_2_FC) versus statistical significance (-log10 *P*-value) for pairwise comparisons (labelled above each plot) for each peak. n-values in corners represent the number of peaks that are enriched (log_2_FC ≥ 1; adjusted *P*-value ≤ 0.05). *P*-values were calculated by the Wald test and corrected for multiple comparisons using FDR. **(C)** Heat map representing number of peaks that are in the trajectories of interest (y-axis) and in each chromatin state (x-axis). **(D)** Heat map of z-scores of the average normalized H3K27ac peak counts for all 4 clusters of interest. Peak sets underwent decision tree clustering based on significant log_2_FC values for associated peaks in each group when compared to VEH-Context. Line graphs in trajectory plots represent significant log_2_FC values for each group in clusters of interest. Count in lower left corner indicates number of peaks. Line plots shown as mean ± SEM. Insets represent differentially enriched active enhancer peaks from each cluster. Normalized counts for each replicate were compared to average normalized count for VEH-Context. **(E)** Gene ontology (GO) analysis of genes associated with H3K27ac peaks. n = 3 biologically independent samples. * *P* < 0.05, ** *P* < 0.01, *** *P* < 0.001

Differential enrichment analysis (Diffbind, DEseq2, data in **Supplemental Table 5**) revealed that CFC in VEH-treated animals led to only marginal changes in H3K27ac enrichment (**Fig. 4B**), in line with the transcriptional data (**Fig. 2A**). Conversely, when CFC occurred in the presence of the HDACi, more then 10,000 and 15,000 regions were significantly down and up-regulated, respectively, indicating that in the presence of HDACi, the behavioral paradigm *per se* can trigger substantial epigenetic changes. Furthermore, the HDACi treatment itself also enriched a significant number of regions – approximately 10,500 regions in both context and CFC treated groups (**Fig. 4B**, **right plots**) – suggesting that both CFC and HDACi treatment alter H3K27ac enrichment. This data is in contrast with the transcriptional results, in which only HDACi treatment, and not the behavioral condition alone, induced transcriptional changes. In addition, while there was equal down and up-regulation of transcription after HDACi treatment (**Fig. 2A, right plots**), we see a higher amount of H3K27ac accumulation after HDACi (**Fig. 4, right plots**).

In order to determine the chromatin state and the corresponding gene for each H3K27ac peak, we used ChromHMM (59) on previously published histone post-translational modifications (PTMs) from bulk hippocampal tissue collected after CFC (30). The entire mouse genome was assigned to one of five chromatin states: Control regions; repressed regions; promoter regions; poised enhancers; and active enhancers (**Supplemental Fig. 12A**). We calculated the state overlap for each peak and assigned the peak to the state that covered the highest proportion (**Supplemental Fig. 12B**). Doing so, 70.5% of bases assigned as active enhancers in ChromHMM were enriched for H3K27ac in our dataset; 44% and 34.9% of bases assigned as poised enhancers promoters, respectively, were also enriched for H3K27ac, while only 2.8% and 2.9% of control regions and repressed regions had H3K27ac peaks (**Supplemental Fig. 12C**).

Next, we performed a decision tree analysis for each chromatin state, focusing on the same four trajectories as before: *primed-active*, *primed-stable*, *primed-silenced* and *non-primed active* (**Fig. 2D-F**). Since active enhancers contained the largest number of peaks (**Fig. 4C**), we chose to analyze this subset of peaks in depth (**Fig. 4D**), but other chromatin states are included in **Supplemental Figure 13**. The *primed-active* cluster for active enhancers was the smallest, containing 179 peaks (**Fig. 4D**). This cluster represented ontologies associated with dendritic locations in the cell (**Fig. 4E**) and included peaks associated with NMDA receptor 2A (*Grin2a*) and Calcium Voltage-Gated Channel Subunit (*Cacna1e*). In the *primed-stable* cluster, there were 378 active enhancers for which H3K27ac was increased after HDACi treatment but not further enriched after CFC (**Fig. 4D**). This cluster included ontologies specific to synaptic locations, and a previously described enhancer of *cFos*, whose regulation by histone acetylation was recently validated by targeted dCas9-p300 manipulations (53). The 1480 enhancers of the *primed-silenced* cluster were also associated with genes that are involved in synaptic assembly and signaling, although noticeably fewer enhancers were associated with IEGs (**Supplemental Table 6**). Finally, the *non-primed active* cluster was the largest and contained 6493 active enhancer peaks, the ontologies for which were also associated with regulation of synaptic signaling. This cluster included enhancers for many genes that are specific for memory and synaptic plasticity. For example, *Fosb*, *Jun*, *Junb* and *JunD*, which are members of the AP-1 complex, known to be involved in neuronal plasticity processes during CFC (41, 42); calcium dependent protein kinases, which are crucial for signaling at glutamatergic synapses (60); and genes in the MAPK/ERK signaling cascade, which regulates H3 acetylation during CFC and helps to establish the stabilization of long-term memory (6, 35, 43).

Taken together, these data show that HDACi-induced H3K27ac enrichment after context or CFC is highly specific to neuronal signaling processes. However, in contrast to the RNA-seq data, the H3K27ac enrichments appear to be most relevant in the *non-primed active* cluster, indicating that it is most responsive to combined HDACi and CFC treatment, which closely resembles the behavioral and electrophysiological results. This is interesting insofar as we would expect changes in acetylation, or our priming step, to be relevant in all HDACi treated groups, but transcriptional activation to be more specific to the paired HDACi and CFC experiments. Thus, to better understand this disconnect between transcriptional activation and acetylation enrichment, we lastly directly compared which genes are both enriched and activated and which genes are only enriched for H3K27ac at enhancer regions.

### Transcriptional activation and H3K27ac accumulation occur at genes involved in synaptic communication

To better understand the relationship between HDACi-induced epigenetic changes and transcriptional activation after CFC, we related H3K27ac accumulation at active enhancers to the expression changes of their associated genes. Doing so, we found that multiple genes underwent a change in their trajectory (**Fig. 5A, B**). The most pronounced trajectory change occurred for genes associated with active enhancers that were in the *non-primed active* cluster in the ChIP dataset, of which 58% changed to being transcriptionally activated by HDACi regardless of whether CFC had occurred or not (*primed-stable*). This indicates that, while CFC was needed to drive their acetylation changes, CFC was no longer required for their transcription changes. These genes were enriched for ontologies including “positive regulation of signal transduction” and “nervous system development” (**Fig. 5C**), whereas ontology analysis for genes that switched between other clusters did not yield any significant hits. Genes in this group included several voltage-gated potassium channels such as *Kcna1*, the transcription factor *Neurod2*, which is crucial for fear learning (61), as well as the IEG and AP-1 complex member, *Jun* (**Fig. 5D**). In addition, the *non-primed active* enhancer to *primed-stable* transcriptional cluster switch was enriched for various genes belonging to the MAPK signaling pathway such as Mapk4, Jun and Rapgef2 (**Fig. 5E and Supplemental Tables 3 and 6**).

**Figure 5.**
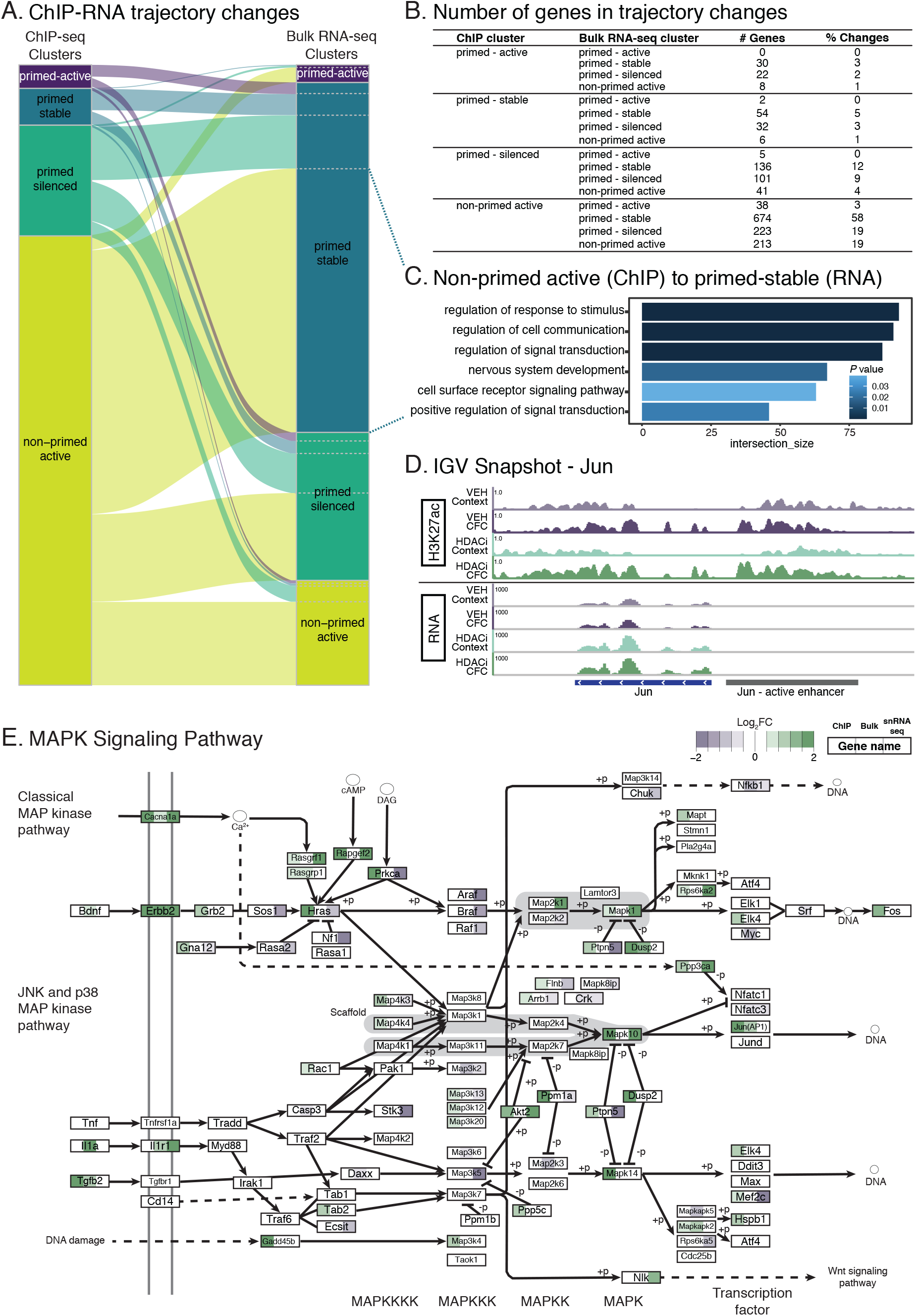
Overlap between HDACi-induced acetylation and transcriptional changes. **(A)** Sankey plot showing the change in trajectory association for the 1585 active-enhancer associated genes present in both the ChIP (left) and bulk RNA-seq (right) and in the trajectory clusters of interest. **(B)** Number of genes changing between the ChIP-seq clusters and the bulk RNA-seq clusters. **(C)** Gene ontologies for the 674 genes in the ChIP non-primed active cluster shifted to the RNA-seq primed-stable cluster. **(D)** Example genome track of the H3K27ac (top 4 tracks) and mRNA expression (bottom 4 tracks) for Jun. Jun’s active enhancer is labelled in grey and the Jun gene is labelled in blue. **(E)** MAPK signaling KEGG pathway visualization. Colors in each box represent significant log_2_FC values for the HDAC-CFC compared to the VEH-CFC comparison in the ChIP (left color), bulk RNA (middle color) and snRNA-sequencing (right color).

When comparing H3K27ac enrichment to the transcriptional activation in single nuclei of the DG after combined HDACi-CFC (**Supplemental Fig. 14A**), we found that only 199 of the 4594 genes were up-regulated in both analyses after combined HDACi-CFC treatment **(Supplemental Fig. 14B**). Despite this being a small subset of the total number of genes, which is likely due to technical differences between the bulk and single nuclei preparations, these genes were relevant to learning and memory in that they included NMDA receptors (*Grin2a* and *Grin2b*), a calcium voltage gated ion channel (*Cacna1e*) and again, members of the MAPK pathway including *Mapk10* and Ras-guanine-nucleotide releasing factor 1(*Rasgrf-1*) (**Fig. 5E**), all of which contribute to glutamatergic synapse communication.

Lastly, when comparing all three datasets together, namely enhancer acetylation, bulk and snRNA-seq transcriptional changes, the MAPK pathway emerged as being predominantly activated (**Fig. 5E**). ERK-mediated MAPK pathway is necessary for memory consolidation (62, 63) and, once activated, ERK phosphorylates protein targets that are implicated in gene transcription, protein synthesis and synaptic plasticity (35), as well as histone acetylation (6). In addition to the MAPK-pathway, 18 genes were increased after combined HDACi-CFC in both the snRNA-seq and bulk-seq and had increased enhancer H3K27ac (**Supplemental Table 7**). Interestingly, two of these genes, autism susceptibility candidate 2 (*Auts2*) and cortactin binding protein 2 (*Cttnbp2*) protect against autism like behavior and impaired object recognition memory (64, 65), while the rest did not seem related to synaptic signaling. Taken together these data suggest that genes involved in synaptic communication and MAPK pathway signaling are epigenetically and transcriptionally activated by HDACi, which suggests these pathways to underlie HDACi-mediated memory enhancement.

## Discussion

In this study, we aimed to determine the mechanisms by which HDACi application facilitates memory formation, and thereby to assess the concept of “cognitive epigenetic priming”. We found that the HDACi CI-994 improves behavioral responses to a subthreshold CFC paradigm (**Fig. 1B**) and following rotarod training (**Supplemental Fig. 5B**), regulated by the hippocampus and striatum respectively. In both behavioral paradigms, CI-994 selectively enhanced unique aspects of synaptic communication within each brain region (**Fig. 1C** and **Supplemental Fig. 5F**) despite these brain areas showing comparably reduced HDAC activity (**Fig. 1E-F**). At the molecular level, HDACi treatment transcriptionally activated distinct gene subsets in each brain region (**Fig. 2**) and between different cell types within the hippocampus (**Fig. 3**). Finally, in DG neurons, HDACi treatment enriched H3K27ac at the enhancers of genes associated with synaptic function (**Fig. 4**), particularly at those involved in MAPK signaling (**Fig. 5**). Together, these findings indicate that CI-994 – although applied systemically – results in brain region, cell type and pathway-specific effects.

As these effects were predominantly observed when HDACi treatment was combined with CFC, but not by either paradigm alone, they support the notion that CI-994 at least partly acts via “cognitive epigenetic priming” (3, 16). This model has been inspired by evidence from cancer research, where HDACi application – inherently devoid of target specificity – improves the efficacy of ongoing cancer treatments, while *per se* having no beneficial effects (17, 18). Analogously, here, we found the HDACi application itself to have minimal effects; but when applied jointly with subthreshold CFC, the HDACi treatment elicited electrophysiological, transcriptional and epigenetic changes that paralleled the improved memory performance.

The brain region-specific electrophysiological effects likely occur because the HDACi treatment reinforces behaviorally relevant cellular pathways per brain area. When paired with CFC, HDACi specifically enhances hippocampal LTP, which is known to underlie contextual fear learning (66–69); whereas when paired with rotarod training, HDACi enhances cortico-striatal PPF, which is known to underlie motor learning (29, 70, 71). This specialization is further supported by the differential transcriptional programs activated in the hippocampus and striatum. While HDACi addition enriched the MAPK pathway in both brain regions irrespective of whether the animals were fear conditioned or only context exposed, the learning and memory-related ERK1 and ERK2 cascade as well as *Bdnf* and *Jun*, which are both involved in MAPK/ERK signaling pathway (34, 53, 72, 73), were only enriched in the hippocampus in combination with contextual learning. This suggests that HDACi generally targets the MAPK pathway but that, when paired with CFC, it leads to a further transcriptional enhancement thereof.

At the epigenetic level, we found a matching correlation between improved contextual memory formation, hippocampal LTP and enhancer H3K27ac enrichment when HDACi treatment was paired with CFC (**Fig. 4**). But even after HDACi treatment alone, we observed H3K27ac-enriched pathways to be mainly associated with synaptic functions. Interestingly, past results have indicated that either HDACi (74) or CFC alone (30) enrich histone acetylation at regions that were already acetylated in baseline conditions. This suggests that the HDACi – although broadly inhibiting HDAC activity – acts by reinforcing acetylated regions, which is likely, given that HDACs are known to be predominantly recruited to and act upon previously activated chromatin regions (75). Furthermore, H3K27ac enrichment also occurred at enhancers of the MAPK pathway (**Fig. 5**), which expands on previous findings linking HDACi treatment to this pathway (6, 76), and testifies to the importance of H3K27ac-induced epigenetic priming for improved memory performance.

At the same time, we observed that H3K27ac changes were not always translated into transcriptional changes (**Fig. 5**), which indicates such gene activation to be independent of H3K27ac priming at this time post-learning. This observation bears striking resemblance to a recent study which described an initial increase in engram enhancer accessibility following CFC, which was not yet paralleled by transcriptional changes, but only after several days post-conditioning (77). In turn, this stipulates that HDACi-induced epigenetic priming might become more important at later stages of memory consolidation. Alternatively, the apparent de-coupling between H3K27ac and the transcriptional changes implies that these changes also rely on other epigenetic modifications. Indeed, several studies have shown that general chromatin rearrangements, a product of combined histone post translational modifications and DNA methylation changes, are necessary for memory formation and occur soon after CFC (30, 41, 42, 77–81).

Given the multifactorial physiological and molecular underpinnings of learning and memory there are several open questions emerging from this study. For example, while we only assessed histone acetylation changes in the DG, we cannot exclude the role of other hippocampal subregions, in particular CA1 (**Supplemental Fig. 4**), to be epigenetically altered by HDACi in response to CFC (67). Another limitation is the possibility that measuring mRNA and histone acetylation changes 1 hour after CFC might be more representative of secondary-wave effects of HDACi application and CFC training, considering that many IEGs, acting as transcription factors themselves, are already up-regulated 30 minutes after CFC (52, 82). Additionally, HDACi effects may reach beyond histone acetylation. For example, HDACi treatment is known to stimulate RNA polymerase II (Pol II) elongation at transcriptionally poised genes by altering PolII acetylation *in vitro* (83, 84). Since many IEGs associated with learning and memory have been found to be in a poised PolII state and are subsequently released in response to neuronal activation in cultures (82), this scenario warrants further investigation *in vivo* as well.

Another interesting observation is the substantial transcriptional down-regulation in response to HDACi (**Fig. 2A)**, which is surprising given that HDACs are members of protein complexes involved in transcriptional silencing (85, 86). Although this phenomenon has been observed in previous studies investigating HDACi treatment alone (74) or when combined with memory extinction (15), it remains to date without definite explanation. Likewise, it remains to be determined whether similar molecular and physiological cascades are triggered by other HDACis or in conditions characterized by impaired cognition.

These open questions notwithstanding, the findings presented here shed light on the mechanisms by which systemic HDACi treatment can lead to specific memory-promoting effects. By enhancing neuronal activity-induced epigenetic and transcriptional cascades, HDACi treatment reaches a high level of target specificity despite being devoid of such specificity *per se.*

## Materials and Methods

### Animals

All procedures, including animal experiments, were handled according to protocols approved by Swiss animal licenses VD2808/2808.1, VD2875/2875.1, VD3169 and VD3413 and according to the standard operating procedures of E-PHY-SCIENCE SAS (ENV/JM/MNO (2077)). Ten-week-old C57BL/6J male mice were purchased from Janvier Labs and allowed an acclimatization and handling period in the EPFL animal house for two weeks before experimentation. All animals were housed in groups of 4-5 animals at 22-25° C on a 12-hour light-dark cycle with food and water ad libitum. Mice were randomly assigned a drug treatment, and experimental conditions were randomly split by cage so that all mice in one cage underwent the same fear conditioning protocol.

### Drug administration

The class I HDAC inhibitor, CI-994 (synthesized at the Broad Institute with a purity of >95% by HPLC analysis)(15), was dissolved in 10% dimethyl sulfoxide (Sigma-Aldrich, D2438), 30% Kolliphor (Sigma-Aldrich, C5135), and 60% 0.9% saline (Braun, 395158) Its vehicle (VEH) solution consisted of all of the above, without CI-994. One hour before contextual fear conditioning, each animal was interperitoneally (i.p.) injected with either 30mg/kg of CI-994 or a corresponding volume of VEH pre-heated to 37°C on a thermomixer. Solutions were made fresh before each experiment and stored at −20°C until use.

### Contextual fear conditioning (CFC)

All behavioral testing was performed between 9AM and 1PM. CFC for behavior, electrophysiology and sequencing experiments consisted of a 3 min habituation to the conditioning chamber (TSE Systems GmbH at EPFL for all molecular experiments; Imetronic (Pessac, France) for electrophysiology experiments) followed by two 1 s foot shocks (0.2mA) with an interval of 29 s and a final 15 s in the chamber. The context groups in all experiments were exposed to the conditioning chamber for the same amount of time with no shocks. The chamber was cleaned with 5% ethanol between each animal.

To measure the effect of CI-994 on fear learning, animals were re-exposed to the chamber for 3 min, 24 h after the initial exposure. Percentage of time spent freezing over the total habituation period was automatically calculated with an infrared beam detection system (MultiConditioning System, TSE Systems GmbH). Freezing was quantified when absence of movement was detected for longer than 1 sec. Animal velocity (average cm/s) and distance travelled (total cm) during the habituation phases were calculated automatically by the TSE system. Changes in anxiety were determined by dividing the conditioning chamber int 36 sections and calculating the percent of total time each animal spent in the inner 16 section (no bordering wall) of the fear conditioning chamber during the initial habituation phase.

For all other molecular experiments, animals were left in their home cage for 1 hour after CFC. Then animals were sacrificed and respective brain regions were manually dissected and immediately frozen on dry ice. Brain regions were stored at −80°C until further processing.

### Rotarod

Motor performance was measured using a Rotarod apparatus (Bioseb, model LE8200). Mice were placed on the rotating rod, and the latency to fall was measured while the speed was accelerating from 4 to 40 rpm. Trials began when mice were placed on the rod and rotation began. Each trial ended, and latency was recorded, when the mouse fell off the rod. Mice were tested for 4 trials with a 1 minute inter-trial interval(87).

### Electrophysiology

One hour after CFC or Rotarod experiments mice were anesthetized with isoflurane and decapitated. Heads were immediately immersed in ice-cold freshly prepared artificial cerebrospinal fluid (aCSF; 124 mM NaCl, 1.3 mM MgSO4, 4 mM KCl, 1.0 mM Na2HPO4, 2.0 mM CaCl2, 26 mM NaHCO3, and 10 mM D-glucose) for at least 2 mins before brain extraction. Acute slices (350 μm thick) were prepared with a vibratome (VT 1000S; Leica Microsystems, Bannockburn, IL) in ice-cold gassed aCSF. Sections were kept at room temperature (RT) for at least 1 h before recording.

For electrophysiological recordings, a single slice was placed in the recording chamber, submerged and continuously super-fused with gassed (95% O2, 5% CO2) aCSF at a constant rate (2 ml/min). fEPSPs were evoked by an electrical stimulation at 0.25 Hz (100μsec duration) in the perforant path, Schaffer collaterals or the cortico-striatal pathway. Downstream extracellular fEPSPs were recorded in the Dentate Gyrus (DG), CA1 and striatum, respectively, using a glass micropipette filled with aCSF. Synaptic transmission input/output (I/O) curves were constructed at the beginning of each experiment to asses basal synaptic transmission. For the I/O, a stimulus ranging from 0 to 100 μA by 10 μA steps was applied and measured every 5 secs. Paired Pulse Facilitation (PPF) was performed to assess short-term plasticity. For PPF, two stimulations were applied and measured at 50, 100, 150, 200, 300 and 400 ms intervals. Stable baseline fEPSPs were recorded by stimulating at 30% maximal field amplitude (single stimulation (0.25 Hz) every 30 secs). The same intensity of stimulation was kept for the reminder of the experiment. After a 10-15 min stabilization, high-frequency stimulation (HFS: 3 trains of 100-Hz stimulation, each train having a 1 sec duration and 2 trains separated by 20 sec) was delivered. Following these conditioning stimuli, a 90 min test period was recorded where responses were again elicited by a single stimulation every 30 sec at the same stimulus intensity. Signals were amplified with an Axopatch 200B amplifier (Molecular Devices, Union City, CA) digitized by a Digidata 1322A interface (Axon Instruments, Molecular Devices, US) and sampled at 10 kHz. Recordings were acquired using Clampex (Molecular Devices) and analyzed with Clampfit (Molecular Devices). Experimenters were blinded to treatment groups for all the experiments.

#### Data Processing

Off-line data analysis of hippocampal and striatal basal synaptic transmission and synaptic plasticity was processed using Clampfit (Molecular Devices). For I/O data, fEPSPs slopes were measured at each intensity of stimulation (from 0 to 100 μA). These slopes were then normalized to the maximal value. Normalized fEPSP slopes were plotted against different intensities of stimulation. PPF measurements were normalized by normalizing the first fEPSP slope to 1 and comparing it with the second fEPSP slope. LTP was measured as percent of baseline fEPSP slope recorded over a 10-min period before HFS delivery. This value was taken as 100% of the excitatory post-synaptic potential slope and all recorded values were normalized to this baseline.

### HDACi assay

Hippocampal and striatal hemispheres, collected 1 hour after CFC consisting of three 2 sec foot shocks (0.8mA), were thawed and homogenized in RIPA buffer (150mM NaCl, 50mM Tris-HCl ph8, 0.1% SDS, 0.5% deoxycolate, 1% NP-40) on ice for 30 min. Proteins were then extracted from the nuclei by adding HDAC buffer (50mM Tris-HCl pH8, 137nM NaCl; 2.7mM KCl; 1mM MgCl2; 1mg/mL BSA) and sonicating at full strength for 15 min (Diagenode, Bioruptor Plus). Protein concentration was measured using a Bradford Assay and normalized so that all assay inputs contained the same amount of protein. Pan-HDAC enzyme activity was determined using the Fluor de Lys HDAC fluorometric activity assay kit (Enzo Life Science, BML-AK500) according to the manufacturer’s protocol. Protein extracts were incubated with the Fluor De Lys Substrate for 30 min and then with the Fluor De Lys Developer for 15 min. Fluorescence intensity (380nm excitation; 510 nm emission) was measured on a the Infinite M200 Pro fluorometric reader (Tecan). Mice treated with VEH and not undergoing fear conditioning were considered as representing baseline HDAC activity (normalized to one-fold). Assays were run in triplicate from 3 independent experiments.

### Western Blots

Animals underwent drug administration and subthreshold CFC as described above. Full hippocampi were dissected and flash frozen 1 h after CFC. Frozen hippocampal hemispheres were cut in half and homogenized and incubated for 30 min on ice in 500μl RIPA buffer (150mM NaCl, 50mM Tris-HCl ph8, 0.1% SDS, 0.5% deoxycolate, 1% NP-40) with 20μl 20x protease inhibitor (Complete mini, EDTA-free, Sigma Aldrich Cat#11836170001). Nuclei were collected by centrifugation (max speed, 20min, 4°C) and cytoplasm (supernatant) was transferred to a new tube. The nuclear pellet was mixed with 50μl 1x Laemmli buffer, sonicated for 10 min at full power and boiled for 10min at 90°C or until samples were no longer viscous. Protein quantifications were performed using a DC assay. For each sample, 10μg protein was added to SDS-PAGE gel (12.5% acrylamide in Resolving Gel and 4.5% in stacking gel) and run at 25A for ~1.5 h. Proteins were then transferred to nitrocellulose membrane for 2 h at 4°C and blocked for 1 h in 5% milk in PBS-Tween20. Primary antibodies (1:2500 H4K12ac (ab46983), 1:500 H3K9ac (ab10812), and 1:5000 H3K27ac (ab4729) in 2% milk + PBS-Tween20) were incubated with the membrane overnight at 4°C (except 1:5000 total H3 (ab1791), incubated for only 30 min at RT). Then membranes were washed 3x in TBS-Tween20 and secondary antibodies (1:10,000 Goat anti-rabbit in 2% milk) for 1 h at RT. Membranes were washed and incubated with chemiluminescent ECL Plus (GE Healthcare, Cat# RPN2232SK) for 5 mins before visualization on the Fusion FX Vilber Laurmat imaging system. Due to similar sizes of histone markers, blotting was done separately and stripped between each antibody.

To quantify chemiluminescence, images were analyzed using “Set Measurements” in ImageJ. For each blot, percent of total luminescence was calculated for each band and normalized to the respective H3 total luminescence. Technical replicates (same samples, 2 western blots) were averaged together for each antibody and per biological replicate (6 replicates per treatment).

### RNA-seq

#### RNA extraction and library preparation

Single frozen hippocampal and striatal hemispheres from four biological replicates were isolated after CFC. Samples were homogenized and total RNA was isolated using Trizol Reagent (Life Technologies) according to the manufacturer’s protocol. RNA was further purified by an on-column DNAse digestion using the RNase-Free DNase Set (Qiagen, Cat# 79254) and two rounds of washes using the RNAeasy Mini Kit (Qiagen, Cat# 74106). Total RNA concentration was determined with the Nanodrop 1000 (v3.8.1, Thermo Fisher).

Libraries were prepared using the TruSeq Stranded mRNA Preparation Kit (Illumina) starting from 900ng of RNA. Libraries were quantified with the dsDNA HS Assay kit (Qubit, Cat# Q32851) and profile analysis was performed using the TapeStation (Agilent, TS4200) D500 Screen Tape System (Agilent, Cat#5067-5588 and 5067-5589). Finally, libraries were multiplexed and sequenced across five lanes on the Illumina HiSeq 4000 (Illumina), yielding 100-bp, paired-end reads, at EPFL’s gene expression core facility.

#### RNA-seq analysis

Truseq adapter sequences were trimmed from raw FASTQ files using bcl2fastq (v2.20.0, Illumina). STAR (v2.6)(88) was used to align FASTQ reads to the mouse mm10 reference genome with annotations downloaded from Ensembl release 93(89). A custom R script was used to count reads mapping to the exonic regions of genes and to define transcript abundance. Reads were only considered if they overlap a single gene region. Differential expression and downstream analysis were performed using DEseq2(90) and custom R scripts. Genes were considered differentially expressed if they had an adjusted p-value ≤ 0.05 and a |log_2_FC| ≥ 0.04. For the trajectory analysis, all experimental groups were compared to samples coming from animals that were treated with the VEH and context paradigm (baseline). Genes were grouped into trajectory pathways using custom-written decision tree clustering in R.

### Nuclear Extraction

Nuclear extraction was performed for both ChIP and single nuclear sequencing experiments. All steps of nuclear extraction were done on ice. First, frozen brain tissue was homogenized in a douncer filled with 6ml Solution D (0.25M Sucrose, 25mM KCl, 5mM MgCl2, 20mM Tris-HCl, pH 7.5). For snRNA-sequencing, 5μg/mL actinomycin D (Sigma, Cat# A4262) was added to Solution D to block transcription induced by disassociation. Samples were centrifuged for 1 min at maximum speed and pellets were resuspended in 4ml Solution D and 2ml Optiprep (Serumwerk Bernburg). Samples were then pelleted by centrifugation for 10 min at 3,200g, and the supernatant was discarded. Optiprep purification was performed a second time.

After the final centrifugation, pellets were resuspended and filtered into 5ml polysterene tubes with filter (75mm) snap-caps (Corning, Cat# 352235). For the ChIP-seq experiments the final resuspension occurred in in PBS-T (0.1% Tween 20) and for snRNA-seq the final resuspension occurred in N-PBS (PBS, 5% BSA, 5μg/mL actinomycinD and 0.2U/μl RNAse inhibitor (Thermo Fisher Scientific, Cat#N8080119)).

### ChIP-seq

#### Nuclear sorting

ChIP-seq was performed on 3 replicates per treatment and each replicate consisted of the pooled dentate gyri from 5 mice. After nuclear extraction (see above), filtered nuclei were cross-linked by incubating with 1% formaldehyde (AppliChem, A08770) for 5 min at RT. Cross-linking was quenched with 125mM glycine (VWR, 101196X) and nuclear structural quality was assessed using an EVOS FL cell imaging system (Life Technologies).

For each sample, approximately 750,000 nuclei were resuspended in 500μl PBS-T (PBS, 0.1% Tween 20). Nuclei were stained with 1:50 Alexa Fluor488 conjugated anti-NeuN antibody (Millipore, MAB377X) for 30 min. Then nuclei were spun down (1250rcf, 4°C, 5 min) and washed in PBS-T (0.1% Tween 20) twice. Finally, nuclei were resuspended and stored in 200μl PBS-T until sorting.

Flow cytometry was performed on the FACSAriaIII (BD Bioscience) by the EPFL Flow Cytometry Core Facility (FCCF). Before sorting, samples were passed through a 26G needle 5 times. Hoechst (1:1000) was mixed into each sample and incubated on ice for 10 mins. Debris was first excluded by gating using forward and side scatter pulse area parameters (FSC-A and SSC-A). Multiplets were then excluded by gating FSC-H vs. FSC-W and SSC-H vs. SSC-W. Single nuclei were sorted by Hoechst intensity, elicited by 405 nm wavelength excitation and measured at 425-475nm (450/50-A). Finally, NeuN+ nuclei were sorted into ice-cold Eppendorf tubes containing 100μl PBS-T.

#### Chromatin immunoprecipitation (ChIP)

After sorting, nuclei were pelleted by centrifugation (4°C, 1250g, 5 min) and lysed by incubating in 750μl RIPA buffer on ice for 10 min. Samples were sonicated on an E220 Focused-ultra-sonicator (Covaris) for 20 min (Peak power = 140W, Duty = 5, Cycle/Burst = 200). Sonication efficiency was measured by decrosslinking 125μl of chromatin in 500μl of TL-Brain Buffer(10mM Tris-HCl pH7.5, 10mM EDTA 200mM NaCl), 50μl of 10% SDS and 1μl of RNAseA (Thermo Fisher, Cat#EN0531) and incubating at 65°C and 650rpm overnight. 10μl recombinant, PCR-grade Proteinase K (Roche, Cat#03115828001) was added and incubated at 45°C and 650 rpm for another hour. DNA was extracted with AcNH4 (100μl of 10M), 20μl glycogen (10μg/μl) and 1ml cold isopropanol and then pelleted by centrifuging at 14000rfc, 4°C for 20min. DNA was further purified in 1ml 70% EtOH and centrifuged (14000rfc, 4°C, 10 min). Sonicated DNA size was assessed on a 1.5% agarose gel.

The rest of the ChIP experiment (beginning from “Protein G Agarose Bead Preparation”) was carried out using the reagents and protocols from the Low Cell ChIP-Seq Kit (Active Motif, 53084). In brief, 400μl of sonicated chromatin was first cleared by incubating with pre-cleared Protein G agarose beads for 2 h on a rotator at 4°C. Half was kept as input for each sample. The other half was immmunoprecipitated overnight at 4°C with 3μl of H3K27ac (Abcam, ab4729). After precipitation, pre-cleared Protein G agarose were added for 3 h, and both input and IP samples were washed following the kit specifications. Cross-linking was reversed by incubating samples with 5μl 5M NaCl and 2μl proteinase K at 65°C, 300rpm overnight. DNA was purified using phenol-chloroform.

#### Library preparation

To prepare libraries for both input and IP samples, the Next Gen DNA Library Kit (Active Motif, Cat# 53216) and Next Gen Indexing Kit (Active Motif, Cat# 53264) were used according to the manufacturer’s instructions. After adaptor ligation, fragments were amplified (1 cycle, 30s at 98°C; 14 cycles, 10s at 98°C, 30s at 60°C and 60s at 68°C) and DNA was cleaned and purified using magnetic SPRI beads (Beckman Coulter, Ca# B23317). Libraries were resuspended in 25μl Low EDTA TE buffer and concentration was measured using a Qubit dsDNA HS Assay Kit. DNA fragments size was determined using a Fragment analyzer (NGS High Sensitivity kit (DNF-474), Agilent). Libraries were sequenced, paired-end, on the Illumina NextSeq 500 at EPFL’s gene expression core facility.

#### ChIP-seq analysis

The Next Gen DNA Library Kit (Active Motif) includes molecular identifiers (MIDs), a 9-base random N sequence that is added with the P5 adaptor, to allow for removal of PCR duplicates from sequencing data. While R1 (75bp) contains the sequence information, R2 (9 bp) contains the MID information. To conserve MID information during mapping, the MID sequence from R2 was appended to the FASTQ header in R1 using a custom R-script. Adapter sequences and low quality regions from R1 were removed using Trimmomatic (v0.38)(91) in single end mode with the following parameters: ILLUMINACLIP:Y2_adapter_seq.fa:0:6:6 SLIDINGWINDOW:10:20 MINLEN:36.

The processed FASTQ file (R1) was then aligned to the mm10 genome using Bowtie2 (v2.3.5)(92) in single-end mode and using default parameters. SAMtools (v1.9) (93) was used to convert SAM files to BAM format and then to sort BAM files. PCR duplicated alignments were removed from the BAM files using a perl script by Active Motif. Finally, multi-mapping and low-quality reads (≥ 40) were removed and BAM files were re-indexed using SAMtools.

Open chromatin peaks were defined using MACS2(94) in broad peak mode. Differentially acetylated regions (DARs) were identified using Diffbind (v2.16.2) (95) and DEseq2(v 1.28.1)(90) with default parameters. Peaks were considered differentially enriched if they had a false discover rate (FDR) ≤ 0.05 and |log_2_FoldChange| ≥ 1.

Since H3K27ac is a marker for both promoters and enhancers, ChromHMM (v1.22)(59) was used to establish a chromatin state model that identified enhancers and promoters. The program was run, allowing for 8 states, on independently published ChIP-sequencing data (30), taken from bulk hippocampal tissue 1 h after CFC. We combined groups to get 5 chromatin states (control regions, repressed regions, promoter regions, poised enhancers and active enhancers) based on the combination of histone marks. This information was aligned with our own peak information to define differentially expressed enhancers and promoters. We assigned enhancers to genes using HOMER (v4.11) annotatePeaks.pl (96). Downstream trajectory analysis was performed (as described in the *RNA-sequencing Analysis* secion) separately for peaks in different chromatins states.

All in-house analysis code can be found at https://github.com/allie-burns/2021_Burns_etal.

### Single-nuclear RNA-seq

#### Library Preparation

For single-nuclear RNA-sequencing (snRNA-seq) animals were treated with either VEH or HDACi and exposed to CFC. For each sample, both hippocampal hemispheres from 5 mice were pooled into two replicates each of VEH and HDACi treated groups. Nuclear extractions were performed as described. Nuclear structural quality was checked using an EVOS cell imaging system and nuclei were counted and diluted to 1,000 nuclei/μl.

#### Library Sequencing

Library constructions were performed using Chromium SingleCell 3’Reagent Kit v3 chemistry (10x Genomics) according to the manufacturers protocol (CG000183 - Rev A). All 4 libraries were pooled and sequenced across 2 NextSeq 500 (v2.5) chips for 75 cycles. FASTQ files were generated using cellranger mkfastq (CellRanger v3.0.1), yielding an R1 length of 28nt and an R2 length of 56 nucleotides.

#### snRNA-seq analysis

To generate single cell feature counts cellranger count (CellRanger v3.0.1) was run to align FASTQ files to the mm10 pre-mRNA genome (created using cellranger mkref (CellRanger v3.0.1)) using the following settings: expect-cells=4000, chemistry = SC3Pv3, r2-length = 56. Downstream analysis was performed using custom R-scripts. Seurat (v4.0.3)(97) was used to calculate quality control metrics. DoubletFinder(98) was used to find and remove doublets and normalization and variance stabilization was done using SCTransform(99). Seurat was then used to perform UMAP and TSNE clustering, to define clusters using molecular identifiers. Differential expression analysis between VEH and HDACi treated groups was performed for each cell type using the logistic regression framework, accounting for replicates, in Seurat’s FindMarkers() command. Augur (51, 100) was used with default commands to calculate perturbation prioritization for each cell type and scProportionsTest (101) to compute cell type composition changes between HDACi and VEH treated samples.

### KEGG pathway visualization

The *Mus musculus* MAPK KEGG pathway (mmu04010) was downloaded from the KEGG PATHWAY Database and drawn using the Bioconductor package, Pathview (102). Differential expression of genes (or enhancers associated with genes) within this pathway are indicated by colors within each box representing a gene: The leftmost color is the log_2_FC value for the active enhancers from the ChIP analysis; the middle color is the log_2_FC for the bulk RNA-seq; and the right most color is the snRNA-seq. The pathway was manually redrawn for visualization purposes and simplified by only plotting MAPK subpathways containing at least one differentially acetylated or transcribed gene.

### Statistics

Statistical details are included in the main text and figure legends, including *P*-values, statistical tests used, ‘n’s for each experiment, and a description of what ‘n’ refers to. Biological replicates refer to biological material from different mice or pools of mice and technical replicates refer to technical repetition using the same material from biological replicates.

## Acknowledgements

We would like to thank all past and current members of the Laboratory of Neuroepigenetics at EPFL for their support and discussion throughout this project, in particular, Paola Arguello and Diego Camacho for their contribution to molecular protocols and analysis. We would also like to thank the EPFL Gene Expression Core Facility (GECF) for their technical assistance with experiment planning and sequencing, the EPFL Flow Cytometry Core Facility (FCCF) for providing the nuclear sorting and the EPFL Center of Phenogenomics (CPG) for ensuring the welfare of the laboratory animals.

