## Supplemental Figures for "The HDAC inhibitor CI-994 acts as a molecular memory aid by facilitating synaptic and intra-cellular communication after learning"

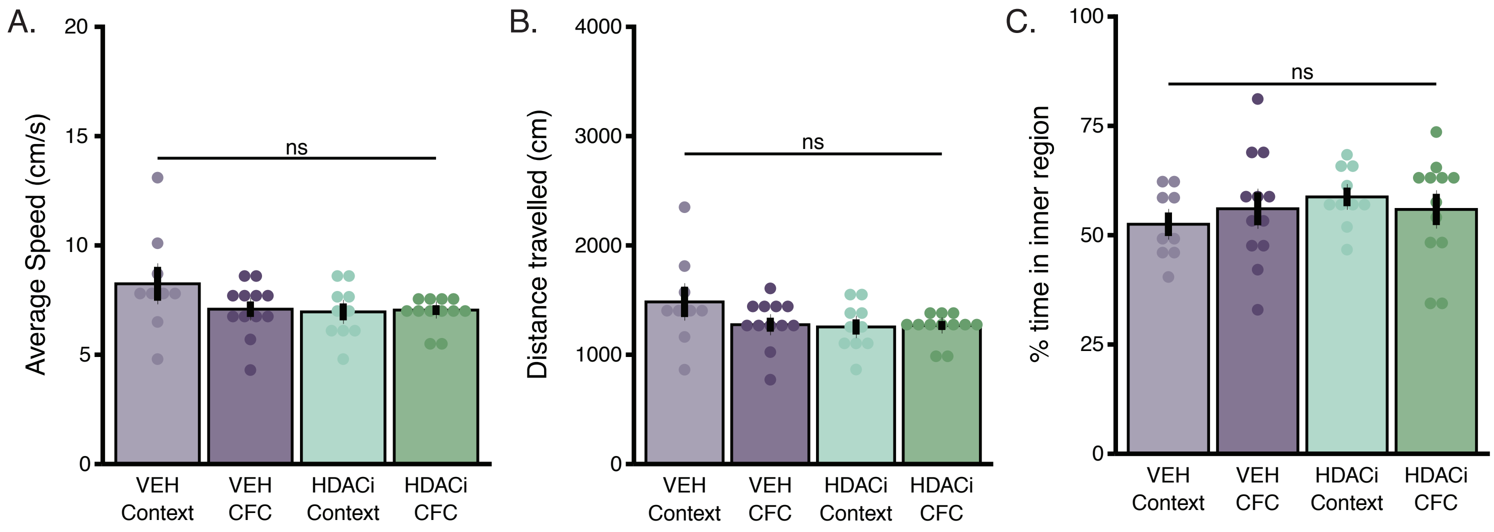


**Supplemental Fig. 1. HDACi treatment does not affect speed, distance travelled or anxiety levels. (A)** Average animal speed (cm/s) during the 3-minute habituation of initial behavioral conditioning was not affected 1 hour after i.p. injection of Vehicle or CI-994. **(B)** Average distance travelled (cm) during the 3-minute habituation of the initial behavioral conditioning was not different 1 hour after i.p. injection of Vehicle or CI-994. **(C)** Time spent in inner regions of the conditioning chamber during the 3 minute habituation of initial conditioning did not change 1 hour after i.p. injection of Vehicle or CI-994. One or two-way ANOVA with Tukey's HSD multiple comparisons test was used for analysis. Graphs represent mean ± SEM. * $P < 0.05$, ** $P < 0.01$, *** $P < 0.001$, ns = not significant. n = 9-12 animals/group

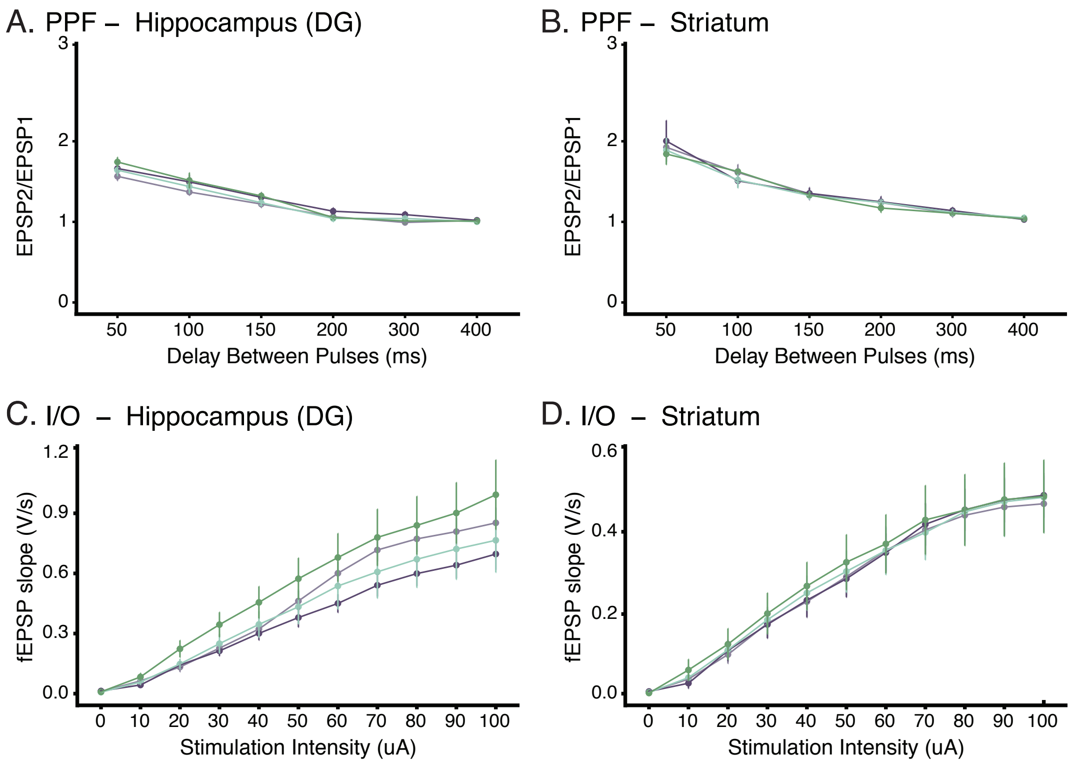


**Supplemental Fig. 2. HDACi does not alter PPF or I/O in the hippocampus or striatum after sub-threshold CFC.** **(A and B)** Paired pulse facilitation (PPF) in the DG **(A)** and striatum **(B)** 1 hour after CFC. **(C and D)** Input/output (I/O) relationship in the DG (C) and striatum **(D)** 1 hour after CFC. Graphs represent mean ± SEM. * $P < 0.05$, ** $P < 0.01$, *** $P < 0.001$, ns = not significant. n = 8 animals/group.

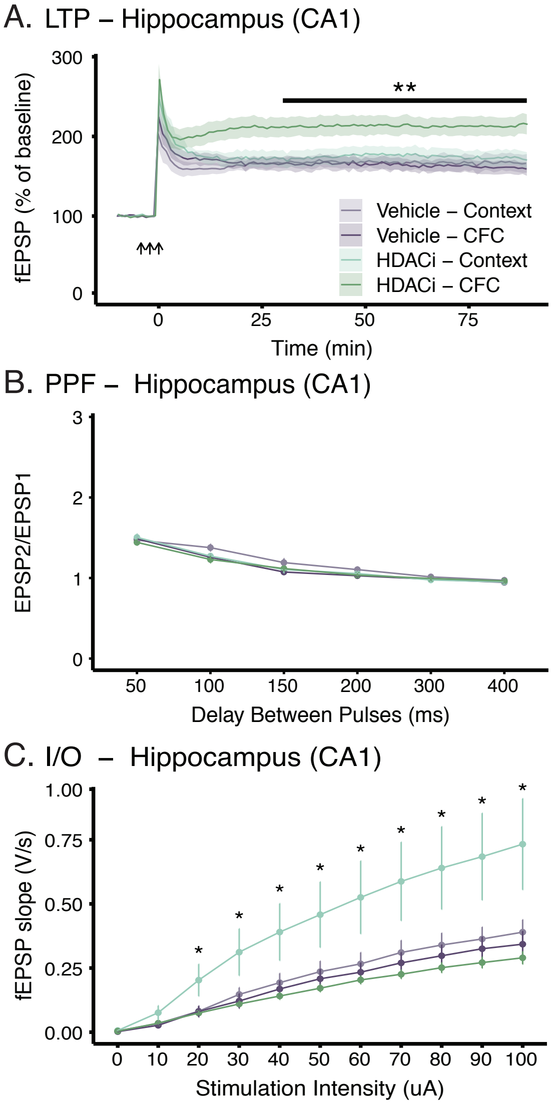


**Supplemental Fig. 3. HDACi combined with sub-threshold CFC enhances LTP in hippocampal area CA1. (A)** HDACi combined with CFC enhanced LTP in response to 3 trains of high frequency stimulation (HFS – arrows) at Schaffer Collaterals of the hippocampal CA1 one hour after initial behavioral conditioning. Statistical differences were calculated for the 30 minutes (end of short-term-potentiation) to 90 minutes (end of recording) for each mouse. **(B)** There were no treatment-induced differences in PPF in the CA1. **(C)** HDACi-Context did have a larger I/O relationship than other groups in the CA1. Graphs represent mean ± SEM. * $P < 0.05$, ** $P < 0.01$, *** $P < 0.001$. n = 8 animals/group.

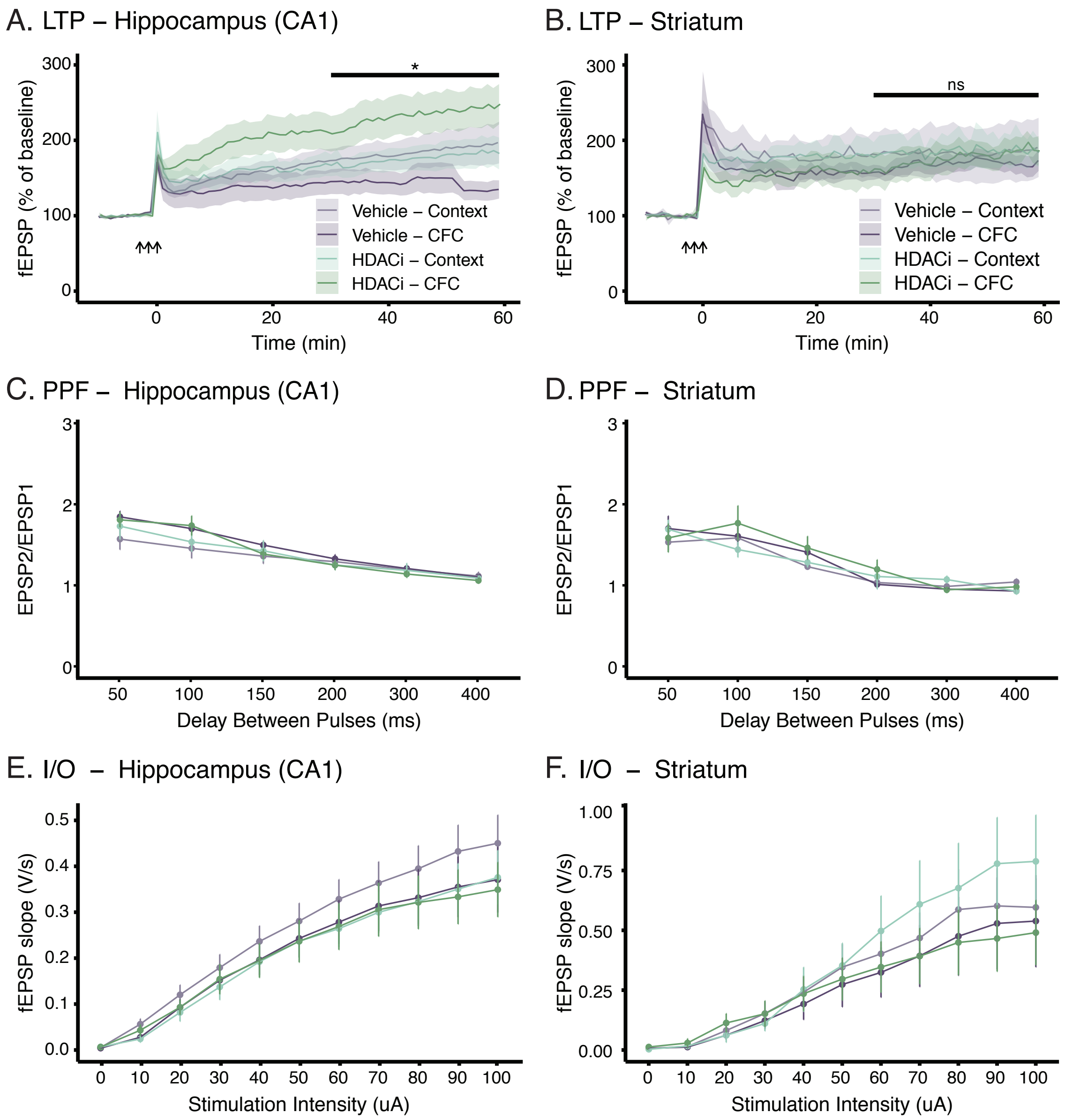


**Supplemental Fig. 4. HDACi combined with strong CFC enhances LTP in CA1 but not in striatum. (A and B)** HDACi combined with strong CFC (3x 0.8mA – 1s) enhanced LTP in response to 3 trains of high frequency stimulation (HFS – arrows) in the Schaffer Collaterals of the hippocampal CA1 **(A)** but not in the cortical-striatal pathway **(B)** one hour after initial behavioral conditioning. Statistical differences were calculated for the 30 minutes (end of short-term-potentiation) to 90 minutes (end of recording) for each mouse. **(C and D)** There are no treatment induced differences in paired pulse facilitation (PPF) in either the CA1 **(C)** or the striatum **(D)** 1 hour after strong CFC. (E and F) There are also no treatment induced differences in the input/output (I/O) relationship in either the CA1 (C) or the striatum **(D)** 1 hour after strong CFC. Graphs represent mean ± SEM. * $P < 0.05$, ** $P < 0.01$, *** $P < 0.001$, ns = not significant. n = 6-10 animals/group.

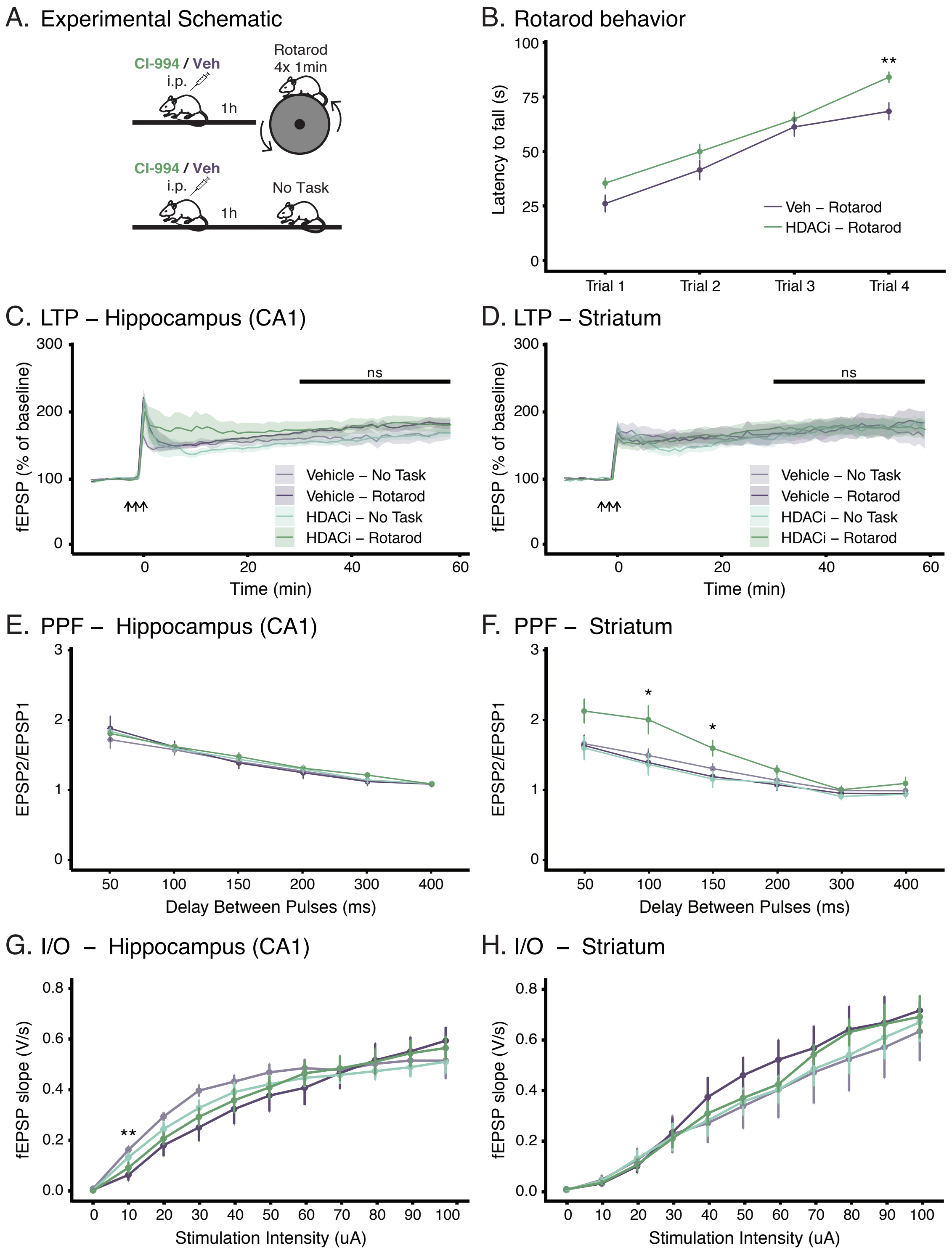


**Supplemental Fig. 5. HDACi enhances PPF in the cortico-striatal pathway after motor learning. (A)** Schematic representation of the striatal specific behavioral paradigm, rotarod training. Animals were i.p. injected with either vehicle or CI-994 (30mg/kg) one hour prior to rotarod training. Rotarod training began by placing each mouse on the rod and accelerating the rod from 4-40rpm in 5 minutes or until the mouse fell off. Mice were tested for 4 trials with 1-minute inter-trial intervals. **(B)** HDACi combined with CFC increases the time to fall off the rotarod after repeated training. **(C and D)** HDACi combined with rotarod training does not alter LTP in response to 3 trains of high frequency stimulation (HFS – arrows) in the Schaffer Collaterals of the hippocampal CA1 **(C)** or in the cortical-striatal pathway **(D)** one hour after final rotarod trials. Statistical differences were calculated for the 30 minutes (end of short-term-potentiation) to 90 minutes (end of recording) for each mouse. **(E and F)** HDACi paired with rotarod training does not lead to any differences in PPF in the Schaffer Collaterals of the CA1 **(E)** but it enhances PPF in the striatum **(F)** after a delay of 100 or 150ms between pulses. **(G and H)** I/O relationships were overall similar in response to HDACi and rotarod training in the Schaffer Collaterals of the CA1 **(G)** or the striatum **(H)**. Graphs represent mean ± SEM. * $P < 0.05$, ** $P < 0.01$, *** $P < 0.001$, ns = not significant. n = 6-10 animals/group.

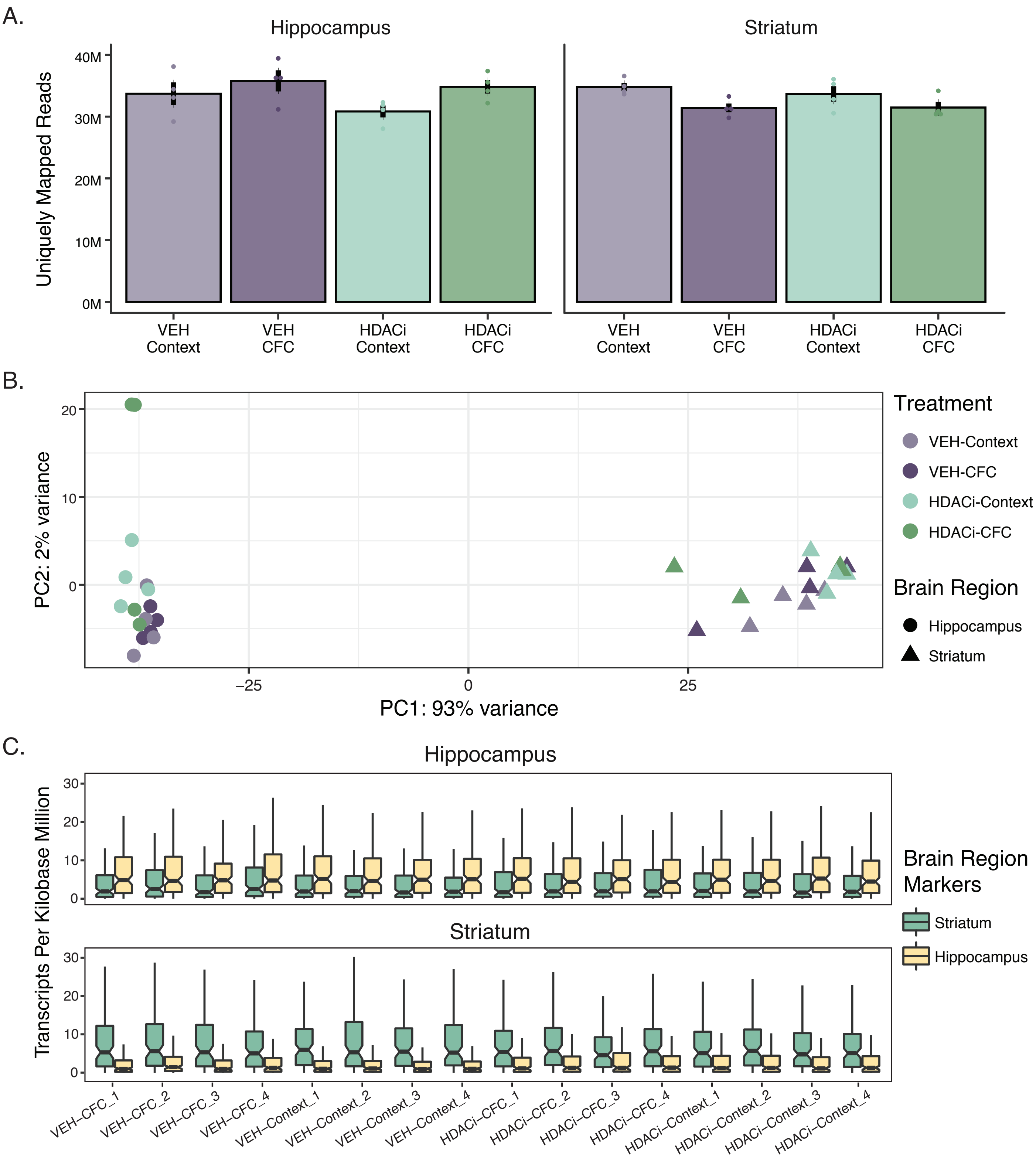


**Supplemental Fig. 6. Alignment statistics for bulk RNA-sequencing. (A)** All libraries contained 28-40M uniquely mapping reads with no treatment group having significantly more reads than any other (One-way ANOVA, $F(7,24) = 2.34$, $P = 0.057$). Graphs represent mean ± SEM. **(B)** Principal component analysis (PCA) showing that 93% of the variance of the top 1000 genes within all 8 libraries comes from differences between brain regions. Only 2% comes from within brain regions. Points are colored by treatment and shape represents brain region. **(C)** Brain region specificity was confirmed by comparing transcripts per kilobase million (TPM) expression of hippocampal and striatal libraries to marker genes from the hippocampus and striatum. The top 250 marker genes for each region compared to all other regions were downloaded from the Allen Brain Atlas. Hippocampal marker genes were mostly highly expressed in libraries from the hippocampus (top), while libraries that were created from striatal tissue were enriched for striatal marker genes (bottom).
n = 4 biologically independent samples.

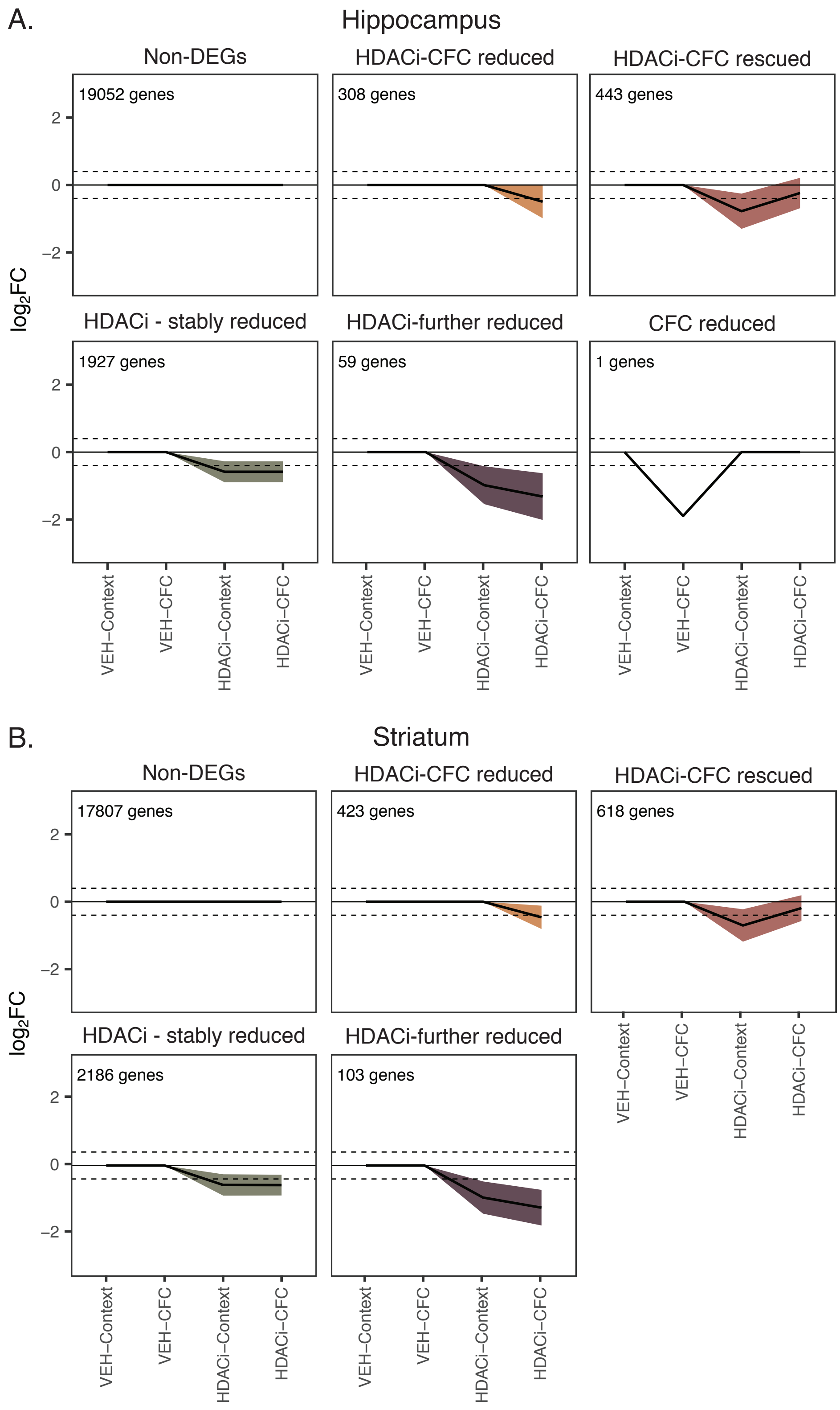


**Supplemental Fig. 7. Clusters representing down-regulated genes in both the hippocampus and the striatum.** Clusters representing non-changed and down-regulated genes in both the hippocampus **(A)** and the striatum **(B).** Most expressed genes in both brain regions were not differentially expressed in any comparison (top left cluster). Both brain regions included similar numbers of genes that were only down-regulated after combined HDACi-CFC (top middle) and genes that were reduced by HDACi-Context treatment and rescued by combined HDACi-CFC (top right). Both brain regions also included genes that were stably reduced by HDACi treatment, regardless of behavioral paradigm (bottom left) and genes that were reduced by HDACi-Context and further reduced by HDACi-CFC (bottom middle). Only the hippocampus contained one gene that was reduced in the VEH-CFC group and rescued by HDACi (bottom right). Line plots represent mean ± SEM

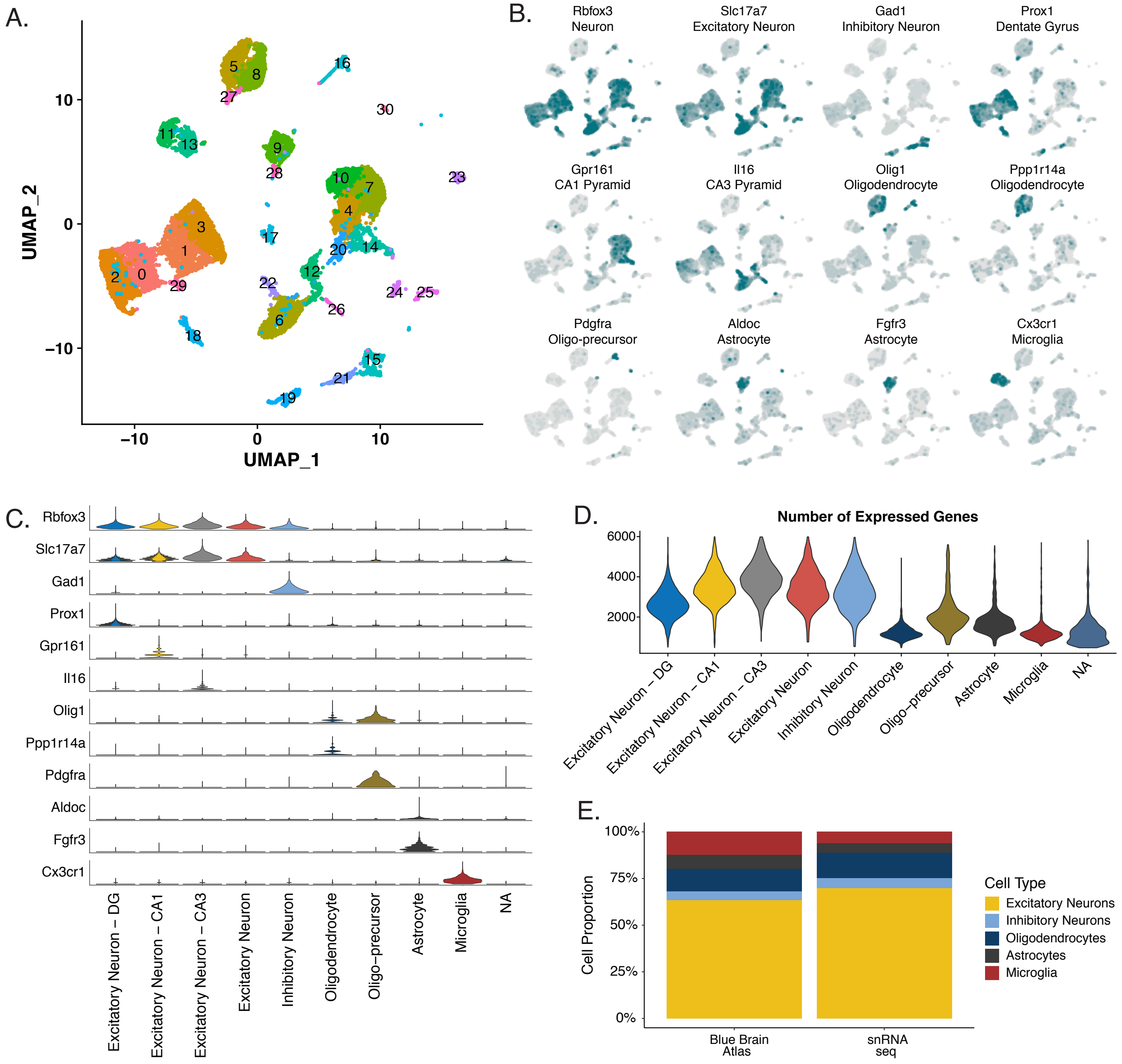


**Supplemental Fig. 8. snRNA-Seq cell type assignments. (A)** snRNA-seq gene count matrices were merged, filtered and normalized using SCTransform and underwent uniform manifold approximation and projection (UMAP) mapping. All nuclei were then clustered based on the k-nearest neighbors. Clusters with fewer than 50 nuclei were removed, yielding 15,339 total nuclei in 30 distinct clusters. **(B)** Clusters were assigned to cell types by overlaying UMI expression for known cell type markers over each cluster. **(C)** Clusters were assigned to cell types (Rbfox3, Slc17a7, Gad1, Olig1, Ppp1r14a, Pdgfra, Aldoc, Fgfr3 and Cx3cr1) and cell locations (Prox1, Gpr161, Il16) based on the markers with the highest UMI expression in each cluster. This analysis yielded 10 distinct cell type and location clusters. **(D)** On average, nuclei assigned to neuronal cell types had more expressed genes than glial cell types (Oligodendrocytes, oligo-precursors, astrocytes, microglia and NA). **(E)** Cell type assignments (right column) were similar to known cell proportions in the hippocampus, taken from the Blue Brain Atlas (left column), irrespective of drug and behavioral treatment. n = 2 biological replicates per group (HDACi-CFC and VEH-CFC).

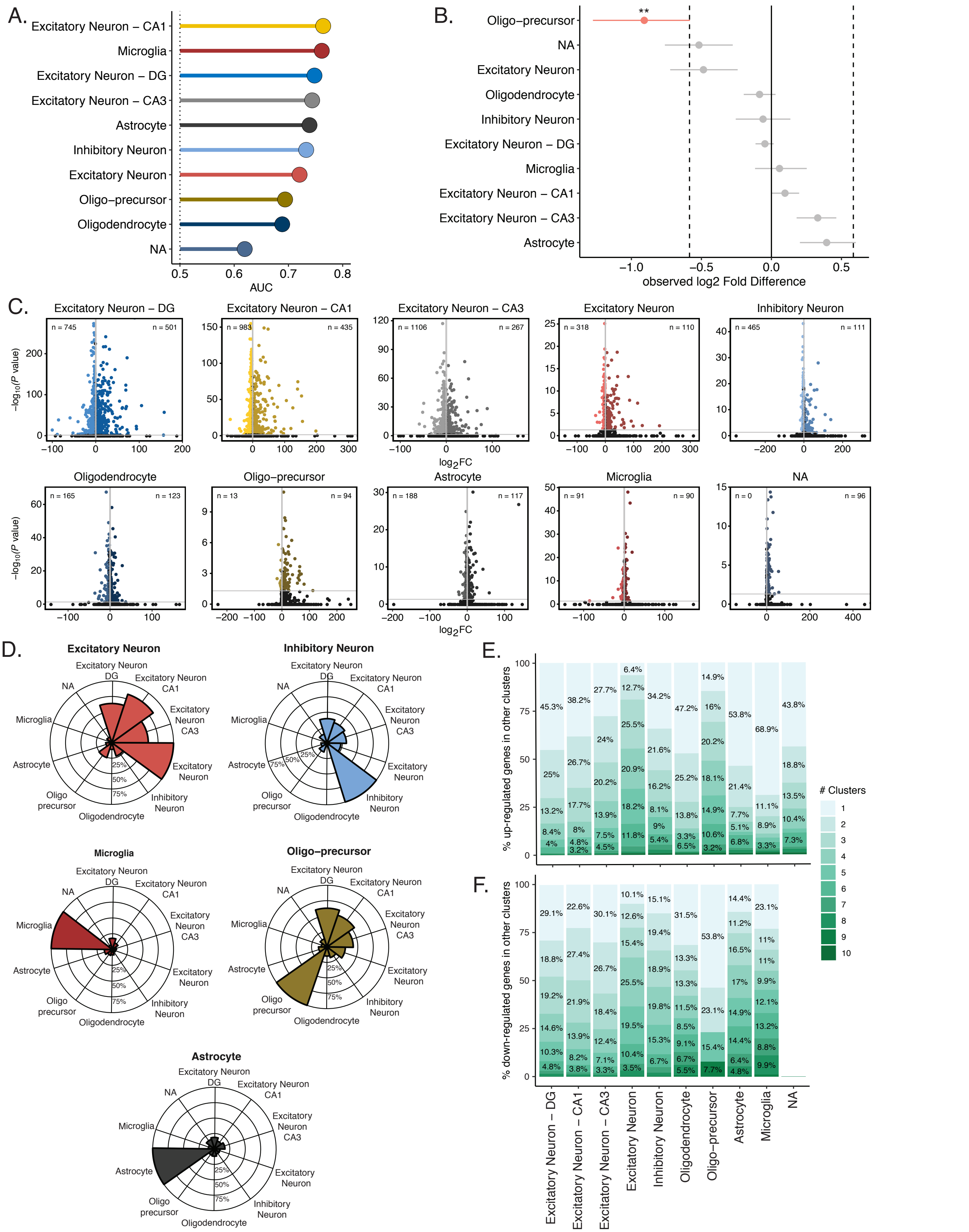


**Supplemental Fig. 9. HDACi perturbs distinct gene sets in different cell types. (A)** Augur analysis. Area under the receiver operating characteristic curve (AUC) is above random chance (0.5) for all cell types. **(B)** Cell type composition permutation test (#permutations = 1000). HDACi only altered relative abundance of cell type by more than 1.5-fold for oligo-precursors. **(C)** Volcano plots showing magnitude of differential expression ($\log_2$FC; $|\log_2\mathrm{FC}| \geq 1$; adjusted p-value ≤ 0.05) versus statistical significance ($-\log_{10}$ P-value) for HDACi-CFC compared to Veh-CFC in each cell type. n-values in corners represent the genes that are up-regulated (right corner) or down-regulated (right corner). **(D)** Overlap of up-regulated genes across cell types not shown in main figure. **(E and F)** Percent of genes found in other clusters for each cell type. Genes that were found in only one cluster are unique to that cell type for up-regulated **(E)** and down-regulated **(F)** genes in each cluster. * $P < 0.05$, ** $P < 0.01$, *** $P < 0.001$, n = 2 biological replicates per group.

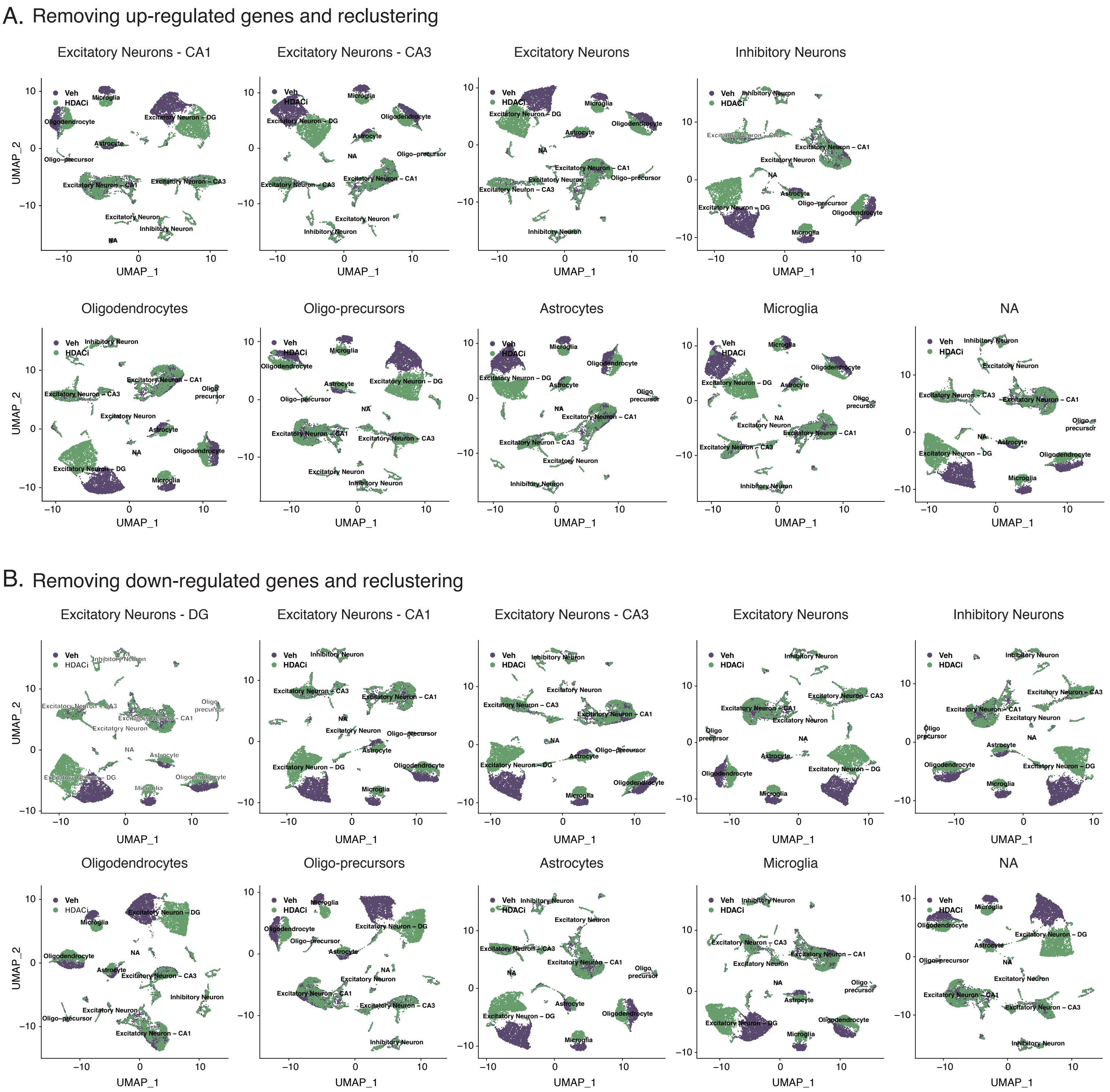


**Supplemental Fig. 10. Removing up- and down-regulated genes and re-running UMAP to determine unique gene sets.** Removing up- **(A)** and down-regulated **(B)** genes and re-running UMAP clustering revealed that HDACi-induced differential expression calculated for each cell type was unique for that cell type. For example, removing up-regulated genes from astrocytes re-merged the astrocyte cluster in the UMAP. Whereas removing down-regulated genes from astrocyte clusters did not.

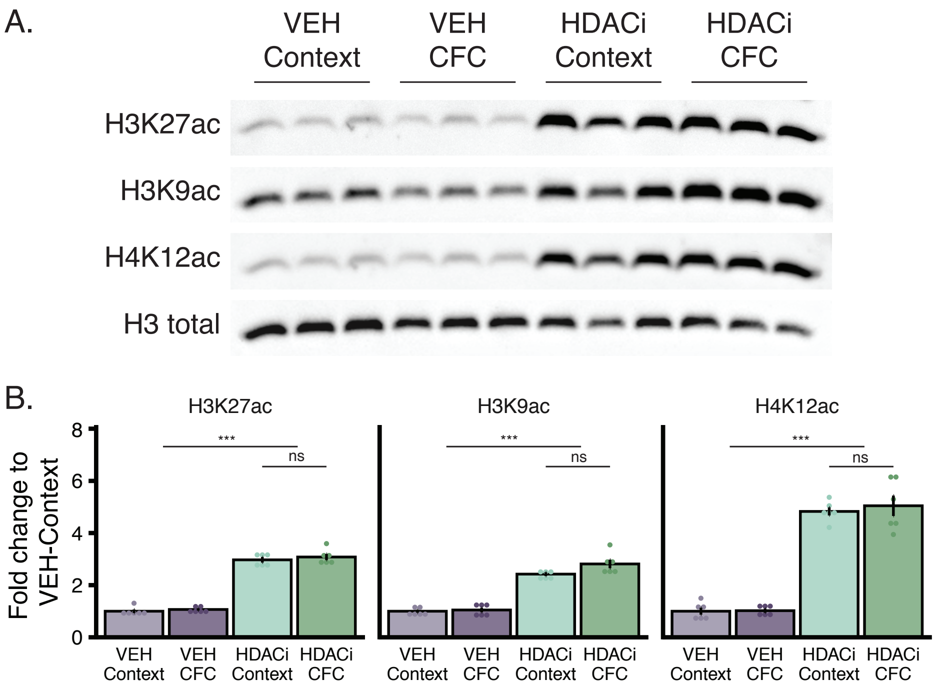


**Supplemental Fig. 11. HDACi enriches H3K27ac, H3K9ac and H4K12ac protein levels regardless of behavioral condition. (A)** Representative Western blot for hippocampal H3K27ac, H3K9ac and H4K27ac including 3 biological replicates for all four treatments and compared to total H3 presence (control). **(B)** Western blot quantification for H3K27ac, H3K9ac and H4K12ac plotted as fold change to the average VEH-Context luminescence indicating that HDACi enriched histone acetylation for all 3 marks shown, regardless of behavioral conditioning. Graphs represent mean ± SEM. n = 6 biological replicates per group, each with 2 technical replicates. Two-way ANOVA, * $P < 0.05$, ** $P < 0.01$, *** $P < 0.001$, ns = not significant.

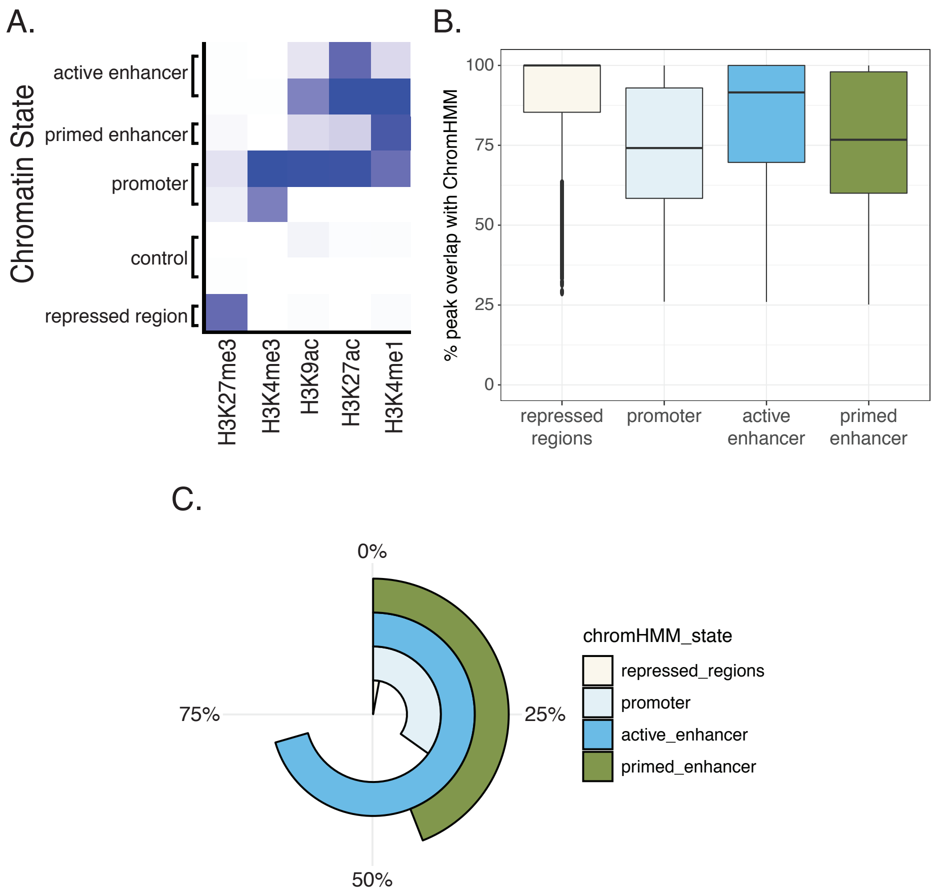


**Supplemental Fig. 12. H3K27ac peaks at enhancer and promoter regions. (A)** Five distinct chromatin states were assigned to the full mm10 genome using previously published histone PTMs and ChromHMM to assign chromatin states. H3K27me3 acts as a repressive marker, H3K4me3 and H3K9ac are promoter markers, H3K27ac is a marker of active enhancers and H3K4me1 is enriched at primed-enhancers. **(B)** Percent of peak overlap with assigned chromatin states shows that most peaks overlap with their assigned state by more than 50%. Graphs shown as box and whisker plots with outliers plotted as points. **(C)** Proportion of each chromatin state (for full genome) enriched by H3K27ac peaks in all four treatments indicates that H3K27ac peaks are enriched at active and primed enhancers and not repressed regions.

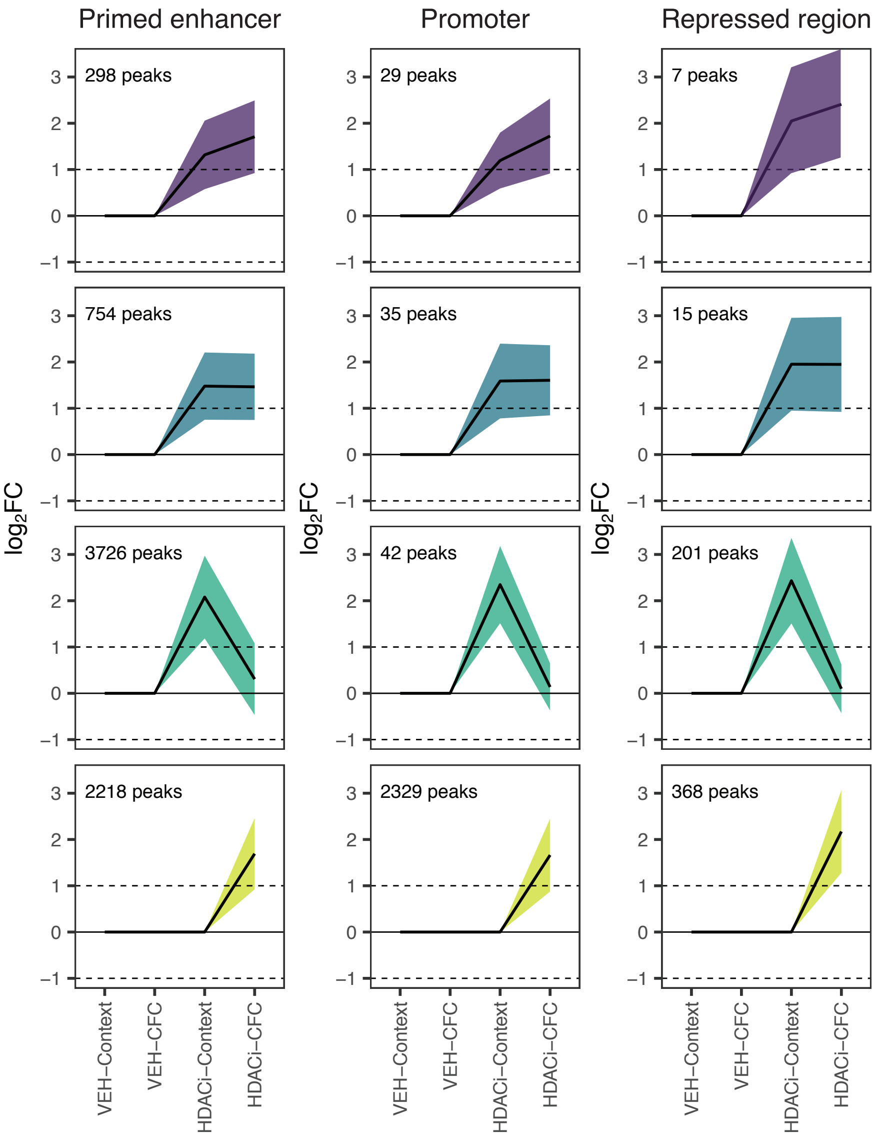


**Supplemental Fig. 13. Trajectory analysis for H3K27ac peaks not associated with active enhancers.** Line graphs in trajectory plots represent significant $log_2FC$ values for each group in clusters of interest for primed enhancers (left), promoters (middle) and repressed regions (right). Count in upper left corner indicates the number of genes in each cluster. Line plots shown as mean ± SEM.

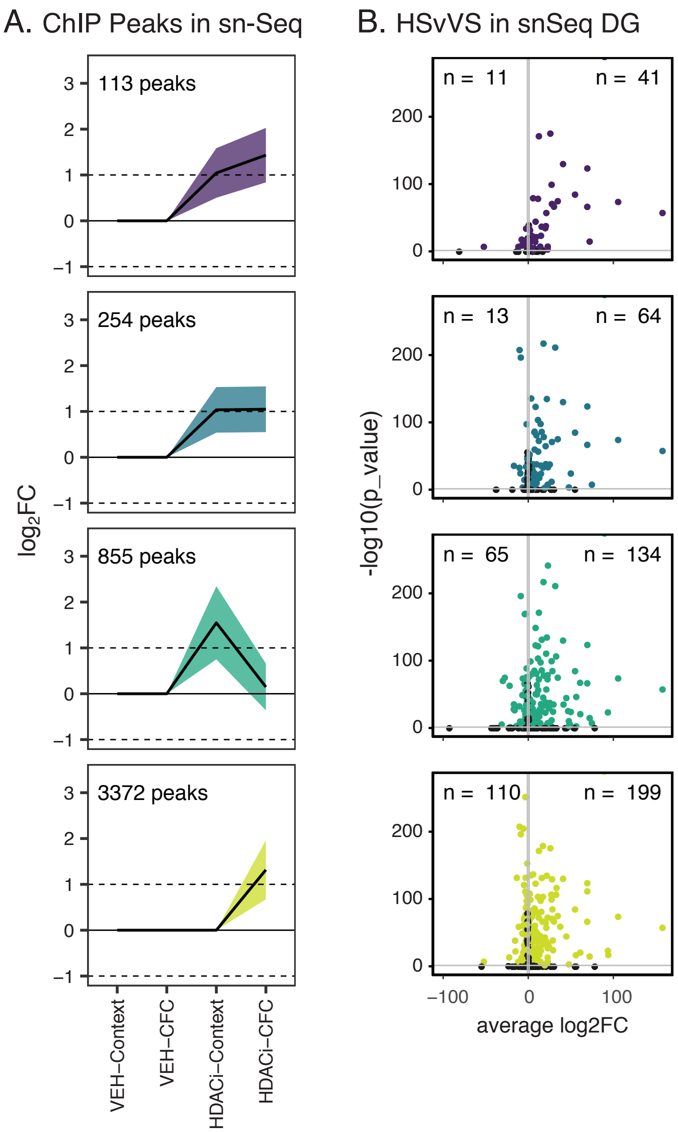


**Supplemental Fig. 14. ChIP to sn-seq comparison. (A)** Line graphs in trajectory plots represent H3K27ac active enhancer peaks that are associated with genes also found in the snRNA-seq analysis. These graphs compare $log_2FC$ values for each group in clusters of interest. Count in lower left corner indicates the number of genes in each cluster. *P*-values for differential expression were calculated using the Wald test in DEseq2 followed by Benjamini-Hochberg multiple comparisons test. Line plots shown as mean ± SEM. **(B)** Volcano plots for genes in the snRNA-seq analysis show that most genes that have increased H3K27ac at active enhancer peaks are also up-regulated in the excitatory neurons of the DG. n-values in corners represent the genes that are up-regulated ($log_2FC \geq 1$; adjusted *P*-value ≤ 0.05).
